# Metabolic constraints drive self-organization of specialized cell groups

**DOI:** 10.1101/573626

**Authors:** Sriram Varahan, Adhish Walvekar, Vaibhhav Sinha, Sandeep Krishna, Sunil Laxman

## Abstract

How phenotypically distinct states in isogenic cell populations appear and stably co-exist remains an unresolved question. We find that within a clonal yeast colony developing in low glucose, cells arrange into metabolically disparate cell groups. Using this system, we model and experimentally identify metabolic constraints sufficient to drive such assembly. Beginning in a gluconeogenic state, cells in a contrary state, exhibiting high pentose phosphate pathway activity, spontaneously appear and proliferate, in a spatially constrained manner. The gluconeogenic cells in the developing colony produce a resource, which we identify as trehalose. At threshold concentrations of trehalose, cells in the new metabolic state emerge and proliferate. A self-organized system establishes, where cells in this new state are sustained by trehalose consumption, which thereby restrains other cells in the trehalose producing, gluconeogenic state. Our work suggests simple physico-chemical principles that determine how isogenic cells spontaneously self-organize into structured assemblies in complimentary, specialized states.

## Introduction

Groups of single, isogenic cells, during the course of development, often form spatially organized, interdependent communities. The emergence of phenotypically heterogeneous, spatially constrained sub-populations of cells is considered a requisite first step towards multicellularity. Here, isogenic cells proliferate and differentiate into phenotypically distinct cells that stably coexist, and organize spatially with distinct patterns and shapes (Newman, 2016; Niklas, 2014). This ability allows groups of cells to maintain orientation, stay together, and specialize in different tasks through the division of labor, all organized with intricate spatial arrangements (Ackermann, 2015; Newman, 2016). In eukaryotic and prokaryotic microbes, the organization into structured, isogenic but phenotypically heterogeneous communities, is widely prevalent, and also reversible (Ackermann, 2015). One well studied example come from the *Dictyostelid* social amoeba, which upon starvation transition from individual protists to collective cellular aggregates that go on to form slime moulds, or fruiting bodies (Bonner, 1949; Du et al., 2015; Kaiser, 1986). Indeed, most microbes show some such complex, heterogeneous behavior, for example in the extensive spatial organization within clonal bacterial biofilms and swarms (Kearns et al., 2004; Kolter, 2007), or in the individuality exhibited in *Escherichia coli* populations (Spudich and Koshland, 1976). Despite its perception as a unicellular microbe, natural isolates of the budding yeast, *Saccharomyces cerevisiae*, also form phenotypically heterogeneous multicellular communities (Cáp et al., 2012; Koschwanez et al., 2011; Palková and Váchová, 2016; Ratcliff et al., 2012; Váchová and Palková, 2018; Veelders et al., 2010; Wloch-Salamon et al., 2017). However, the rules governing the emergence and maintenance of new phenotypic states within isogenic cell populations remain unclear.

Current studies emphasize genetic and epigenetic changes that are required to maintain phenotypic heterogeneity within a cell population (Ackermann, 2015; Sneppen et al., 2015). In particular, cells can produce adhesion molecules to bring themselves together (Halfmann et al., 2012; Halme et al., 2004; Octavio et al., 2009; Váchová and Palková, 2018), or support possible co-dependencies within the populations. However, these studies do not provide an underlying biochemical logic to explain how distinct, specialized cell states can emerge and persist in the first place. This is more so in isogenic, and therefore putatively identical, cells in seemingly uniform environments. Contrastingly, a common theme in all described examples is the requirement of some ‘metabolic stress’ or nutrient availability that is necessary for the emergence of phenotypic heterogeneity and spatial organization, often in the form of metabolically inter-dependent cells (Ackermann, 2015; Campbell et al., 2016; Cáp et al., 2012; Johnson et al., 2012; Liu et al., 2015). Experimentally, systems-engineered metabolic dependencies between non-isogenic cells can result in interdependent populations that constitute mixed communities (Campbell et al., 2016, 2015; Embree et al., 2015; Wintermute and Silver, 2010). These findings suggest that biochemical constraints derived from metabolism may determine the nature of phenotypic heterogeneity, and the spatial organization of cells within the population. It is therefore important to understand what these constraints are, and how they can explain the self-organization of genetically identical cells into distinct states. Here, using clonal yeast cells, we experimentally and theoretically show how metabolic constraints imposed on a population of isogenic cells can determine shared resource production, utilization, and the spontaneous emergence of cells exhibiting a counter-intuitive metabolic state. These constraints drive the overall self-organization of genetically identical cells into specialized, spatially ordered communities.

## Results

### Cells within *S. cerevisiae* colonies exhibit ordered metabolic specialization

Using a well-studied *S. cerevisiae* isolate as a model (Reynolds and Fink, 2001), we established a simple system to study the formation of a clonal colony with irregular morphology. On 2% agar plates containing a complex rich medium with low glucose, *S. cerevisiae* forms rugose colonies with distinct architecture, after ∼5-6 days (Fig 1A). Such colonies do not form in typical, high (1-2%) glucose medium (Figure 1A). Thus, as previously shown (Granek and Magwene, 2010; Reynolds and Fink, 2001), glucose limitation (with other nutrients being non-limiting) drives this complex colony architecture formation. Currently, the description of such colonies is restricted to this external rugose morphology. With such a description, as observed in Figure 1A, the colony surface has an internal circle and some radial streaks near the periphery. We carried out a more detailed observation of entire colonies under a microscopic bright-field (using a 4x lens). This unexpectedly revealed distinct internal patterning, and apparent spatial organization of cells within (Figure 1B). As categorized purely based on visual optical density (‘dark’ or ‘light’), regions between the colony center and periphery had optically dense (dark) networks spanning the circumference of the colony, interspersed with optically rare regions. In contrast, the periphery of the colony appeared entirely light (Figure 1B). Based simply on these optical traits alone, we categorized cells present in these regions of the colony as dark cells and light cells (Figure 1B). At this point, our description is visual and qualitative, and does not imply any other difference in the cells in either region. However, this visual description is both robust and simple, and we use this nomenclature for the remainder of this manuscript.

**Figure 1:**
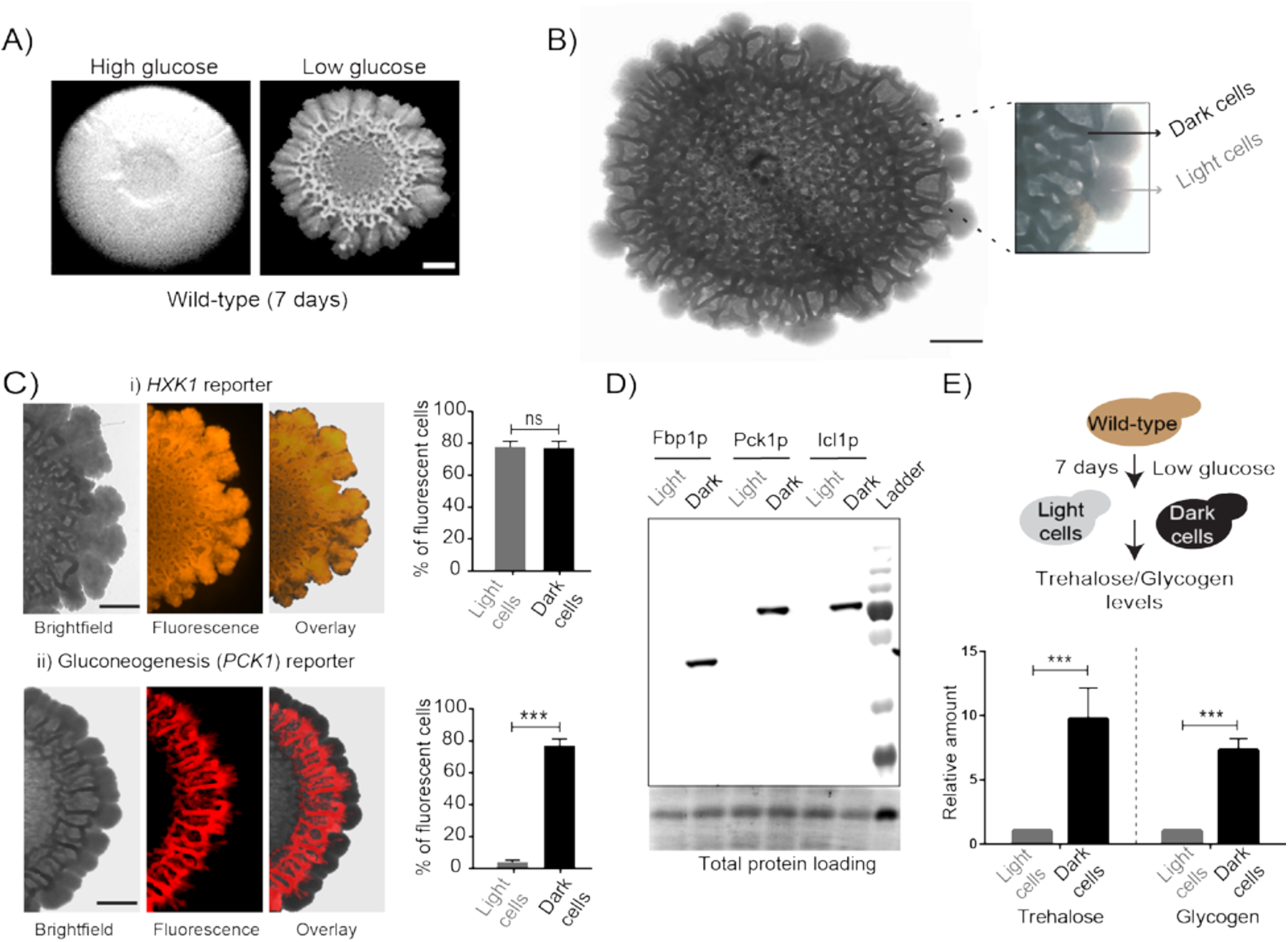
Cells within S. cerevisiae colonies exhibit ordered metabolic specialization. A) Low glucose is required for rugose colonies to develop. The panel shows the morphology of mature yeast colonies in rich medium, with supplemented glucose as the sole variable. Scale bar: 2 mm. B) Reconstructed bright-field images of a mature wild-type colony. Within the colony, a network of dark and bright regions is clearly visible, as classified based purely on optical density. We classify the cells in the dark region as dark cells, and in the peripheral light region as light cells. Scale bar: 2 mm. C) Spatial distribution of mCherry fluorescence across a colony, indicating the activity of (i) a reporter for hexokinase (*HXK1*) activity, or (ii) a gluconeogenesis dependent reporter (*PCK1*), in two different colonies. The percentage of fluorescent cells (in isolated light and dark cells from the respective colonies) were also estimated by flow cytometry, and is shown as bar graphs. Scale bar: 2 mm. Also see Figures S1A and S1B for more information. D) Western blot based detection of proteins involved in gluconeogenesis (Fbp1p and Pck1p), or associated with increased gluconeogenic activity (Icl1p), in isolated dark or light cells. The blot is representative of three biological replicates. E) Comparative steady-state amounts of trehalose and glycogen (as gluconeogenesis end point metabolites), in light and dark cells.

Since these structured colonies form only in glucose-limited conditions, we hypothesized that dissecting the expected metabolic requirements during glucose limitation might reveal drivers of this internal organization. To explore this hypothesis, we first designed visual indicators for hallmarks of yeast cell growth in low glucose. Under glucose-limited conditions, all cells would be expected to have constitutively high expression of the high-affinity hexokinase (Hxk1p) (Lobo and Maitra, 1977; Rodríguez et al., 2001). Further, during glucose-limited growth, all cells are expected to carry out extensive gluconeogenesis, as the default metabolic state (Broach, 2012; Haarasilta and Oura, 1975; Yin et al., 2000). We therefore designed two different fluorescent reporters, one dependent on *HXK1* expression (mCherry under the *HXK1* gene promoter), and the second on *PCK1* expression as an indicator of gluconeogenic activity (mCherry under the *PCK1* gene promoter) (Figure S1A). These reporters were expressed in cells, seeded to develop into colonies, and the expression levels of these were observed in the mature, rugose colony (5-6 days). Expectedly, the *HXK1* dependent reporter showed constitutive, high expression in all cells across the entire colony (Figure 1C). Contrastingly, only the dark cells exhibited high gluconeogenesis reporter activity, which was entirely lacking in the light cells (Figure 1C). To quantify this, cells were dissected out from dark or light regions respectively (under the light microscope, using a fine needle), and the percentage of fluorescent cells in each region was measured using flow cytometry. Based on flow cytometric readouts, >80% of the dark cells showed strong fluorescence for the gluconeogenic reporter, while the light cells (∼97%) were non-fluorescent (Figure 1C, Figure S1B). This suggested a spatial restriction of gluconeogenic activity to within only the dark cell region. We therefore directly estimated native protein amounts of enzymes associated with gluconeogenesis (Pck1-phosphoenolpyruvate carboxykinase, Fbp1-fructose-1,6-bisphosphatase, and Icl1-Isocitrate lyase which is part of the glyoxylate shunt active during gluconeogenesis) in isolated light cells and dark cells. Only the dark cells showed expression of gluconeogenic enzymes (Figure 1D). Finally, we measured steady-state amounts of trehalose and glycogen within dark and light cells, using these metabolites as unambiguous readouts of the biochemical outputs of gluconeogenesis (François et al., n.d.). We observed that the dark cells had substantially higher amounts of both trehalose and glycogen (Figure 1E), indicating greater gluconeogenic activity in these cells. Collectively, these results reveal that intracellular gluconeogenic activity is spatially restricted to specific regions, resulting in a distinct pattern of metabolically specialized zones within the colony.

### Cells organize into spatially restricted, contrary metabolic states within the colony

In the given nutrient conditions of low glucose, gluconeogenesis is expected to be a constitutive metabolic process, essential for all cells. This can therefore be considered as a necessary, permitted metabolic state in this condition. Paradoxically, in these mature colonies, gluconeogenic activity was spatially restricted to only within the dark cell region, with no discernible gluconeogenic activity in the cells located in the light region. This absence of gluconeogenic activity in these light cells, concomitant with a constitutively high level of hexokinase activity, therefore poses a .biochemical paradox. What might the metabolic state of these light cells be? To address this, we first compared the ability of freshly isolated light and dark cells to proliferate in both gluconeogenic (low glucose), and non-gluconeogenic (high glucose) growth conditions. Isolated light cells and dark cells were inoculated either into a medium where gluconeogenesis is essential (ethanol+glycerol as a sole carbon source), or in high glucose medium where cells rely on high glycolytic and pentose phosphate pathway (PPP) activity, and initial growth was monitored. Here, cells that had been growing in high glucose were used as a control. Expectedly, the dark cells proliferated with minimal lag when transferred to the gluconeogenic medium (Figure 2A). However, the light cells showed an extended lag phase in this medium (Figure 2A). Conversely, light cells grew robustly and with minimum lab when transferred to the high glucose medium, as compared to the dark cells (Figure 2A). Counter-intuitively, this indicated that despite being in a low-glucose environment, the metabolic state of light cells was suited for growth in high glucose.

**Figure 2:**
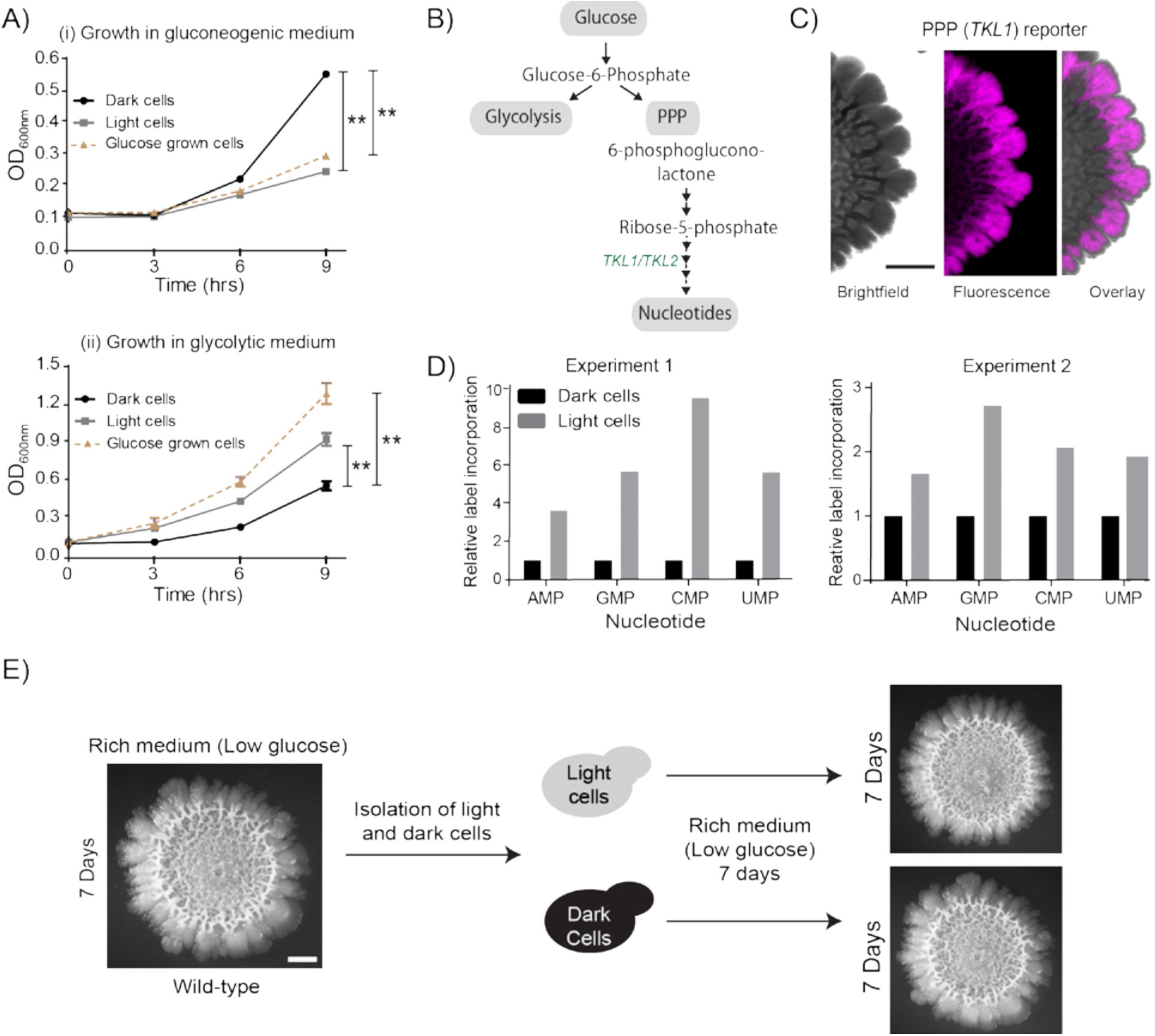
Cells organize into spatially restricted, contrary metabolic states within the colony. A) Comparative immediate growth of isolated light cells and dark cells, transferred to a ‘gluconeogenic medium’ (2% ethanol as carbon source), or a ‘glycolytic medium’ (2% glucose as carbon source), based on increased absorbance (OD_600_) in culture. Wild-type cells growing in liquid medium (2% glucose) in log phase (i.e. in a glycolytic state) were used as controls for growth comparison. B) A schematic showing metabolic flow in glycolysis and the pentose phosphate pathway (PPP), and also illustrating the synthesis of nucleotides (dependent upon pentose phosphate pathway). *TKL1* controls an important step in the PPP, and is strongly induced during high PPP flux. C) Spatial distribution of mCherry fluorescence across a colony, based on the activity of a PPP-dependent reporter. Scale bar: 2 mm. Also see Figure S1A. D) Comparative metabolic-flux based analysis comparing ^15^N incorporation into newly synthesized nucleotides, in dark and light cells. Also see S2, and Materials and Methods. E) Light cells and dark cells isolated from a 7-day old wild-type complex colony re-form indistinguishable mature colonies when re-seeded onto fresh agar plates, and allowed to develop for 7 days. Scale bar = 2mm.

In the presence of glucose, yeast cells typically show high glycolytic and PPP activities, as part of the Crabtree (analogous to the Warburg) effect (Crabtree, 1929; De Deken, 1966) (Figure 2B). Therefore, if the light cells in the colony were indeed behaving as though present in glucose-replete conditions, they should exhibit high PPP activity. To test this, we designed a fluorescent PPP-activity reporter (mCherry under the control of the transketolase 1 (*TKL1*) (Walfridsson et al., 1995) gene promoter, Figure S1A), and monitored reporter activity across the mature colony. Indeed, only the light cells exhibited high PPP-reporter activity (Figure 2C). Next, we directly assessed if the biochemical outputs corresponding to high PPP activity were high in the light cells. High nucleotide synthesis is a canonical consequence of enhanced PPP activity (Nelson and Cox, 2013). The carbon backbone (ribose-5-phosphate) of newly synthesized nucleotides is derived from the PPP, while the nitrogen backbone comes from amino acids (Nelson and Cox, 2013) (Figure 2B, Table 2). We devised a metabolic flux-based experimental approach to assess *de novo* nucleotide biosynthesis in light and dark cells, as an end-point readout indicative of high PPP activity. Light and dark cells, isolated from colonies were pulsed with a ^15^N-label (ammonium sulfate+aspartate), and incorporation of this label into nucleotides was measured by liquid chromatography/mass spectrometry (LC/MS/MS). Light cells had higher flux into nucleotide biosynthesis, compared to the dark cells (Figure 2D, Table 2). Taken together, light cells exhibit the metabolic hallmarks of cells growing in glucose-replete conditions, including increased PPP activity, and increased nucleotide biosynthesis. Thus, in the spatially organized colony, the light cells and dark cells have contrary metabolic states. This is despite the prediction that the gluconeogenic state, exhibited by the dark cells, is the plausible metabolic state in the given growth conditions.

**Table 1.**
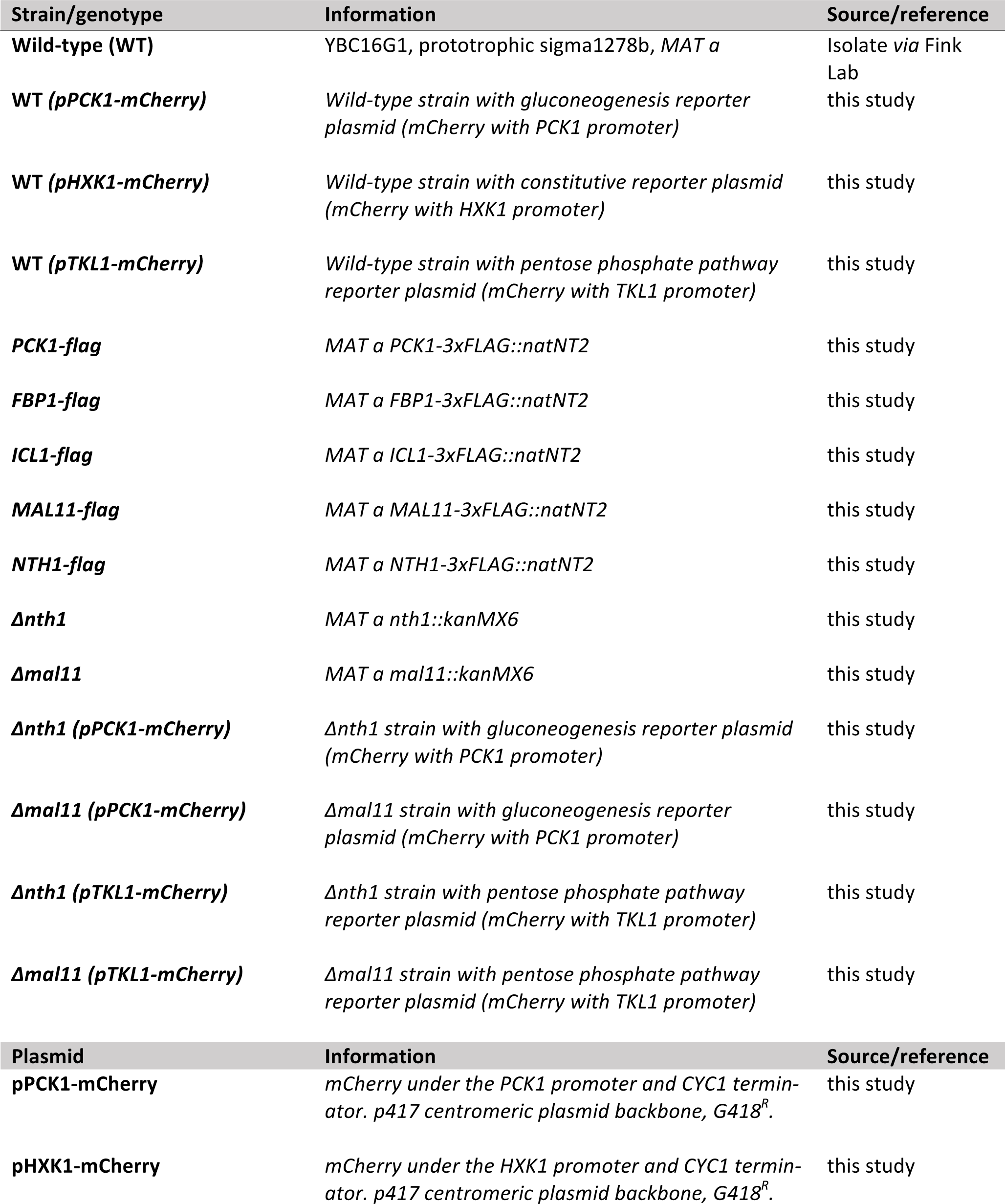
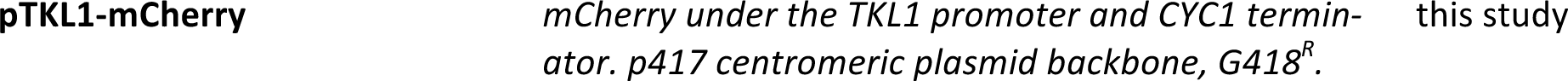
Strains and plasmids used in this study.

**Table 2.**
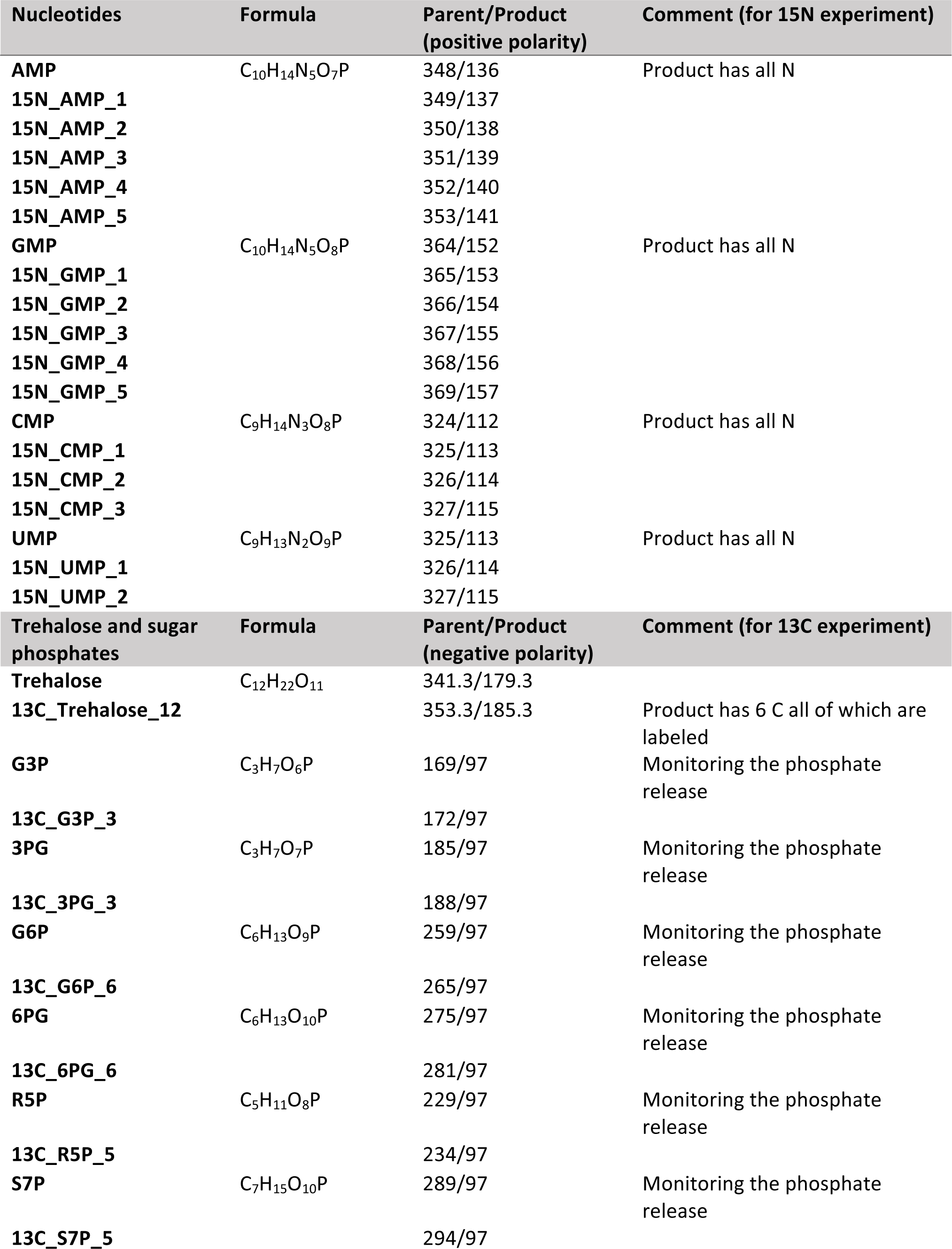
Mass transitions used for LC-MS/MS experiments.

Notably, the light cells or dark cells, when isolated and reseeded as a new colony, both develop into indistinguishable, complex colonies (Figure 2E). This reiterates that these phenotypic differences between the light and dark cells are fully reversible, and do not require genetic changes. Collectively, these data reveal that cells organize into spatially separated, metabolically specialized regions. Within these regions, cells exhibit complimentary metabolic states, one of which is counter-intuitive and cannot obviously be sustained given the external nutrient environment.

### A mathematical model suggests constraints for the emergence and organization of cells in complimentary metabolic states

What determines the emergence and spatial organization of a group of cells, in these contrary metabolic states? Particularly, what explains the emergence and proliferation of the light cells, which exhibit a counter intuitive metabolic state, while maintaining some subset of cells in the dark state? To address this, we built a simple mathematical model. This model incorporates simple processes derived from our current experimental data, to simulate the formation of a colony of ‘light’ and ‘dark’ cells. The model was intentionally coarse-grained, since its purpose was only to find a minimal, biologically consistent combination of processes that is sufficient to produce the overall spatial structure and composition of cell states observed in the colonies. The intention behind the model was not to decipher all possible molecular details that explain this phenomenon. The model should sufficiently account for both the emergence of light cells, as well as their spatial organization with dark cells. Therefore, such a model could suggest constraints that determine the emergence of light cells, which can then be experimentally tested.

While building our model, we included a range of processes that must be considered, based on our experimental data thus far (Figure 3A). This includes the dark cells switching to a light state, the production of some resource by dark cells, diffusion parameters for the resource, consumption of the resource, and rates of cell division (Figure 3A). Next, we constructed a two-dimensional square grid of ‘locations’ for groups of cells within the colony (Figure 3B), where each location is either empty or occupied by a group of ∼100 cells (also see Materials and Methods for full details). Note: we intentionally coarse-grain the grid (for computational simplicity to simulate colony sizes comparable to real colonies) by approximating that the locations either consist of all light or all dark cells. At each time step (12 min of real time), the processes shown in Figure 3A are executed all across the spatial grid using the outlined algorithm (Figure 3C). In this algorithm, (i) all cells consume available nutrients (present in saturating amounts), while glucose concentrations are negligible, (ii) dark cells grow and divide in the given conditions, (iii) dark cells produce a resource/resources as a consequence of their existing gluconeogenic state, (iv) this resource diffuses around the grid and is freely available, (v) dark cells switch to the light state if sufficient resource is present at their location, and lastly, (vi) the resource can sustain the light state cells, which can expand if there is an empty location in the neighborhood, and if the resource is not there the light cells can switch back to dark. All processes occur at specified rates, allowing for stochasticity. Finally, this existence of a shared resource is surmised because, logically the emergence of light cells from dark can happen only if the local nutrient environment enables a switch to the new metabolic state.

**Figure 3:**
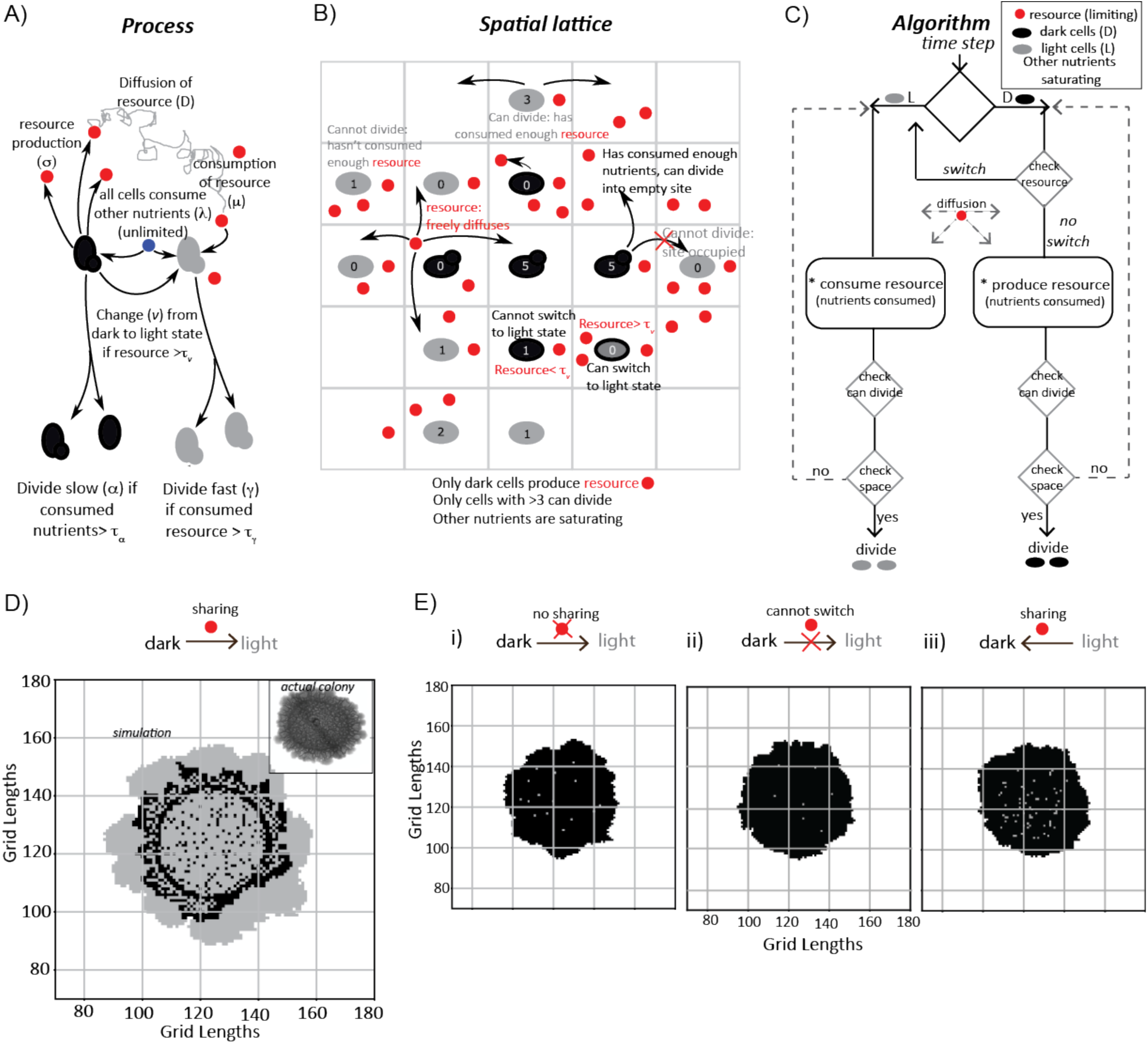
A mathematical model suggests constraints for the emergence and organization of cells in complimentary metabolic states. A) Processes, based on experimental data, incorporated into developing a simple mathematical model to simulate colony development. The dark and light cells are appropriately colored, and the parameters incorporated are resource production (s), diffusion parameters for the resource (D), consumption of the resource (m), and fast or slow rates of division (a or g), based on resource or amino acid consumption. B) The spatial distribution of cells is reduced to a grid like lattice within the model, to allow coarse graining of the location of cells across a colony. The rules for cell division and expansion incorporate the ability to consume existing nutrients in the medium, produce a resource and/or consume a produced resource, and a threshold amount of resource build up before utilization. C) A flow-chart of the algorithm used in the mathematical model. The decision making process in the algorithm, incorporating all the elements described in panels (A) and (B) is illustrated. Also see Figure S2 and Materials and Methods. D) A simulation of the development of a wild-type colony, based on the default model developed. The inset shows an image of a real wild-type colony, which has developed for an equivalent time (∼6 days). Also Figure S2 and Movie S1. E) A simulation of colony development using the model, where key parameters have been altered. (i) The sharing of a produced resource is restricted. (ii) The ability to switch from a dark to a light state is restricted. (iii) Light cells produce a resource taken up by dark cells is included. Note that in all three scenarios the colony size remains small, and fairly static. Also see Figure S3 and Movies S2, S3 and S4.

In each simulation, empty grids are seeded with 1257 occupied locations, with 95-99% of the cells in the dark state. After ∼750 time steps (corresponding to ∼6 days) a simulated wild-type colony looks typically as shown in Figure 4D (also see Figure S2 and Video S1). Strikingly, the simulated spatial organization (Figure 3D, S2 and Video S1) recapitulates most features of a real colony (Figure 3D). These are: at the edge of the initial circular inoculation is a ring of dark cells, the outermost part of the colony is made up of outcrops of light cells, and from this ring of dark cells emanate clusters of dark cells penetrating into the outcrops of light cells. This is despite the simplicity of the rules in the model, including its flattening into 2D. In the simulation, for the first 40-45 time steps, the colony remains small and predominantly dark, while the resource builds up. Then, dark cells start to switch to light. When this happens within the bulk of the colony, these light cells have restricted division due to spatial constraints. Around 100 to 150 time steps later, light cells emerge at the perimeter of the colony, and then rapidly divide and expand (Figure 3D, Video S1). In order to test if the processes of Figure 3A are all required for this behavior, we examined three comprehensive control scenarios: (i) dark cells do not produce a resource (for growth light cells depend only on amino acids, or other pre-supplied resources), (ii) dark cells cannot switch to the light state, and (iii) light cells produce a resource that is needed by dark cells to grow (a straw-man). None of these cases produces the wild-type spatial organization, over a wide range of parameter values (Figure 3E, Figure S3 and Videos S2-4).

**Figure 4:**
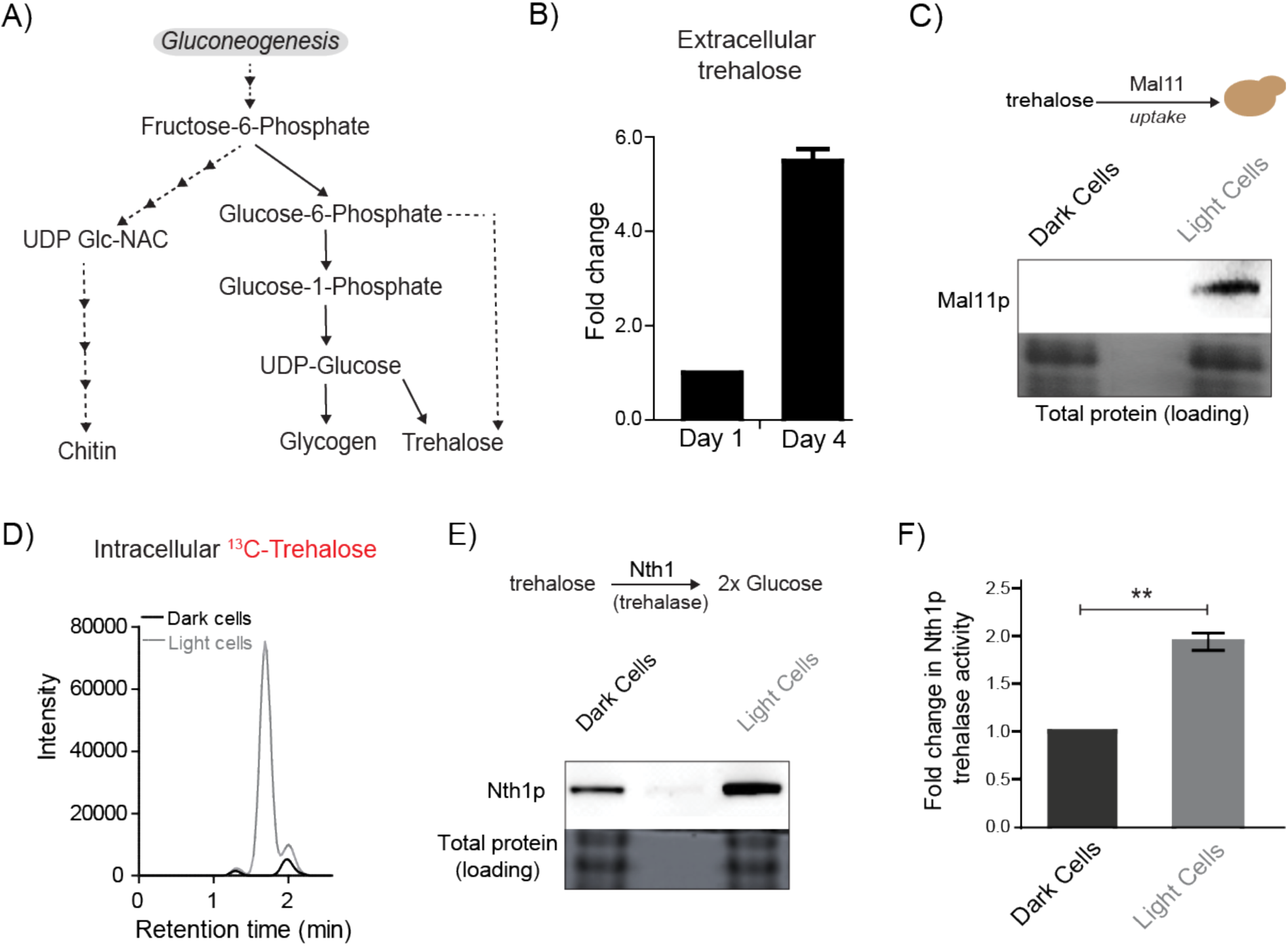
Trehalose satisfies criteria to be the metabolic resource determining the emergence of light cells. A) A schematic illustrating the metabolic intermediates and different end-point metabolites of gluconeogenesis. B) Extracellular amounts of trehalose measured from developing wild-type colonies. Entire colonies were isolated, and only exogenous trehalose estimated, at the respective days. C) Comparative protein amounts of Mal11, a major transporter of trehalose in *S. cerevisiae*, in light and dark cells, as measured using a Western blot is shown. The blot is representative of 3 independent experiments. D) Estimates of the relative ability of light and dark cells to uptake trehalose is shown. ^13^C Trehalose was exogenously added to light and dark cells, and intracellular amounts of the same are shown (as intensity of the MS/MS peak corresponding to ^13^C-trehalose). E) Comparative amounts of Nth1, the major intracellular trehalase enzyme in *S. cerevisiae*, in light and dark cells, as measured using a Western blot is shown. The blot is representative of 3 independent experiments. F) *in vitro* neutral trehalase activity present in lysed light or dark cells is shown. The data represents mean±SD, n=3 (p<0.01).

Summarizing, this simple model successfully recaptures the general features of the spatial patterning and organization of real colonies. This includes the overall general architecture, and spatial organization of light and dark cells. Two simple take-home points emerge from this model. First, the model requires that dark (gluconeogenic) cells produce a resource that is needed by dark cells to switch to the light state. Second, a resource produced by the dark cells is required to sustain the light state. Collectively, in our model, these metabolic constraints are sufficient to determine the overall spatial organization of metabolically distinct, specialized cells.

### Trehalose satisfies criteria to be the metabolic resource determining the emergence of light cells

Does any gluconeogenic metabolite(s) determine the organization of these cells, consistent with these requirements suggested by experimental and modeled data? Such a metabolite must logically satisfy the following three criteria. First, this resource should be available in the extracellular environment (i.e. released by cells), second, cells must selectively take up this resource, and third, the resource should be metabolized to produce glucose/a glucose-like product capable of fueling a glycolytic and PPP-active state. Further, if this were a ‘controlling resource’, preventing the uptake and utilization of the resource should prevent the emergence and proliferation of only the light cells. To identify such a candidate metabolite, we considered all outputs of gluconeogenesis: the storage carbohydrates/sugars glycogen and trehalose, the polysaccharides of the cell wall (chitin, mannans, glycans), and glycoproteins (Figure 4A) (Jules et al., 2008; Kayikci and Nielsen, 2015). The large size of glycogen, chitins, and complex glycosylated proteins, the lack of known cellular machinery for their uptake, and the difficulty in efficiently breaking them down make them all unlikely candidates to be the controlling resource. Contrastingly, trehalose is a small, non-reducing disaccharide composed of two glucose molecules. Trehalose has been observed in the extracellular environment (Parrou et al., 2005), and has uptake transporters in yeast (Jules et al., 2008; Stambuk et al., 1998). Further it can, when rapidly liquidated to two glucose molecules, fuel glycolysis and re-entry into the cell division cycle (Laporte et al., 2011; Shi et al., 2010a; Shi and Tu, 2013). This presented trehalose an excellent putative candidate metabolite that controlled the emergence of cells in the light state. To test this possibility, we first measured extracellular trehalose in colonies. Free trehalose was readily detectable in the extracellular environment of these colonies (Figure 4B). To test if trehalose could be differentially transported into either light or dark cells, we first estimated amounts of a primary trehalose transporter, Mal11 (Stambuk et al., 1998) in these cells. Mal11 protein amounts were substantially higher in the light cells compared to the dark cells (Figure 4C). To unambiguously directly estimate trehalose uptake, we isolated light and dark cells from a mature colony, and exogenously added ^13^C-trehalose. We then measured intracellular levels of labeled trehalose in these cells (by LC/MS/MS) (Table 2). The light cells rapidly accumulated ^13^C-trehalose (Figure 4D), while the dark cells did not, suggesting robust, preferential uptake of extracellular trehalose. Finally, we estimated the ability of light and dark cells to break-down and utilize trehalose. For this, we first measured the expression of the predominant neutral trehalase in yeast (Nth1) (Jules et al., 2008), in the light and dark cells. Light cells had substantially higher Nth1 amounts than the dark cells (Figure 4E). We also measured enzymatic activity for Nth1 (*in vitro*, using cell lysates), and found that the light cells had ∼2 fold higher *in vitro* enzymatic activity, compared to the dark cells (Figure 4F). Collectively, these data suggested that the light cells were uniquely able to preferentially take up more trehalose, break it down to glucose, to potentially utilize it to sustain a metabolic state with high PPP activity.

### Trehalose uptake and utilization determines the existence of light cells

Since these data suggested that trehalose uptake and utilization would be preferentially high in the light cells, we directly tested this using a quantitative metabolic flux based approach. To the isolated light and dark cells, ^13^C-trehalose was externally provided, and metabolites extracted from the respective cells. The intracellular amounts of ^13^C -labeled glycolytic and PPP intermediates were subsequently measured using LC/MS/MS (Figure 5A, Figure S4A, Table 2). ^13^C –labeled glucose-6-phosphate (which enters both glycolysis and the PPP), the glycolytic intermediates glyceraldehyde-3-phosphate and 3-phosphoglycerate, and the PPP intermediates 6-phosphogluconate, ribulose-5-phosphate and sedoheptulose-7-phosphate all rapidly accumulated exclusively in the light cells (Figure 5A, Figure S4A). Since the labeled carbon comes directly from trehalose, these data indicate both the breakdown of trehalose to glucose, and the subsequent utilization of glucose for these pathways. Thus, external trehalose is preferentially taken up by the light cells, and utilized to fuel the complimentary metabolic state of the light cells, with high glycolytic and PPP activity. Finally, we tested if the sharing and differential utilization of trehalose determined both the emergence and proliferation of light cells. An explicit prediction is made both in our model and our hypothesis based on these experimental data. This is: preventing sharing and utilization of trehalose should prevent cells from switching to the light state. To test this prediction, we generated strains lacking *NTH1* (which cannot liquidate trehalose), and *MAL11* (which will have reduced trehalose uptake), allowed them to develop into mature colonies, and compared the amounts of light cells in each. Compared to wild-type colonies, cells lacking the major trehalose uptake transporter (*Δmal11*) formed colonies with very few light cells, and cells lacking trehalase (*Δnth1*) had nearly no detectable light cells in the colonies formed, based on brightfield observations (Figure 5B). This was more directly observed in colonies of cells with these genetic backgrounds, using the expression of the fluorescent PPP reporter. Here, almost no PPP reporter activity was observed in the *Δnth1* cells (Figure 5D). As controls, we ensured that there were no defects in the expression of the PPP reporters in cells from these genetic backgrounds. Also note: while Mal11 shows a high affinity for trehalose, *S. cerevisiae* has other sugar transporters with reduced affinity for any disaccharide. Therefore, cells lacking *MAL11* may take up trehalose with lower efficiency, but these cells are still capable of breaking down trehalose. Correspondingly, the percentage of dark, highly gluconeogenic cells (as determined using the gluconeogenesis reporter) was proportionately higher in the *Δmal11* (∼73%), and *Δnth1* (∼80%) colonies compared to the wild-type colony (∼65%) (Figure 5C). Thus, controlling the uptake and utilization of the resource directly regulates the emergence of cells in the light state. Collectively, these data demonstrate that trehalose is the shared gluconeogenic resource that determines the emergence and persistence of light cells within the structured colony.

**Figure 5:**
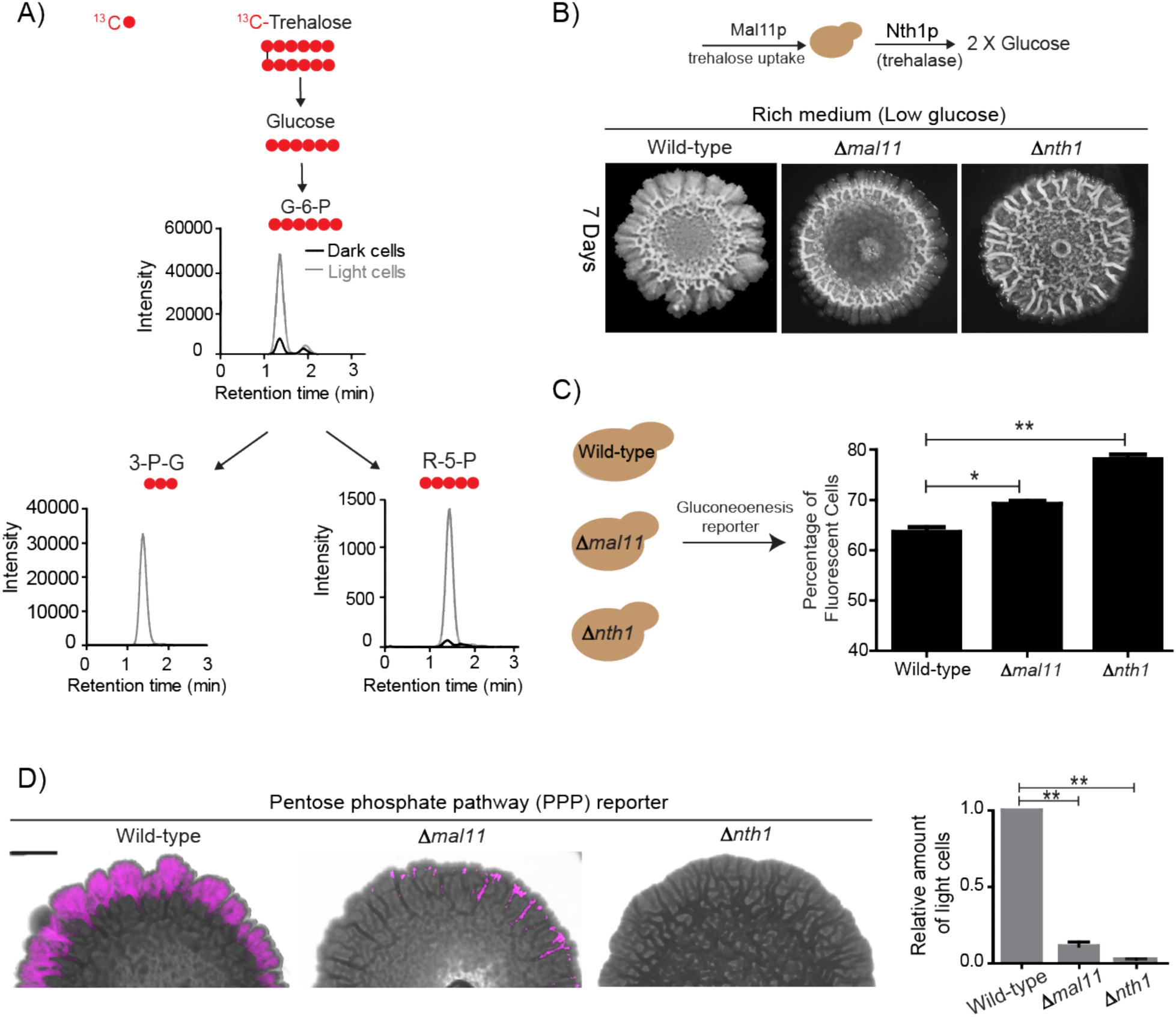
Trehalose uptake and utilization determines the existence of light cells. A) Estimation of trehalose uptake and breakdown/utilization in light and dark cells. LC-MS/MS based metabolite analysis, using exogenously added ^13^C Trehalose, to compare breakdown and utilization of ^13^C Trehalose for glycolysis and the PPP, in light and dark cells. The red circles represent ^13^C labeled carbon atoms. Data for ^13^C labeled glycolytic and PPP intermediates (derived from trehalose) are shown. The data presented is from a single flux experiment, which was repeated independently (with different colonies) twice. Also see Figure S4. B) Comparative development of wild-type colonies with colonies lacking the major trehalose transporter (*Dmal11*), or the intracellular neutral trehalase (*Dnth1*). Colonies are shown after 7-days of development. Scale bar: 2 mm. C) Estimate of the percentage of gluconeogenic cells in wild-type, *Δmal11* and *Δnth1* (strains that cannot up-take or breakdown trehalose). This was based on quantifying the expression of the gluconeogenesis reporter plasmid (pPCK1-mCherry), expressed in all these cells. Cells from the entire colony were isolated and percentage of fluorescent cells (i.e. cells expressing the gluconeogenic reporter) in each colony was calculated by analyzing the samples by flow cytometry. Error bar represents standard error of mean (SEM) * = P<0.05, ** = P<0.01. D) Visualization (right panel) and quantification (left bar graphs) of light cells in wild-type, *Δmal11*, or *Δnth1* cells, based on fluorescence emission dependent upon the PPP reporter activity. The quantification is based on FACS data. Scale bar: 2 mm.

### A resource threshold effect controls cooperative switching of cells to the light state

Our experimental data showing the organization of dark and light cells was obtained from ∼5-6-day old, mature colonies. However, in our simulations of the temporal development of the colony, we observed that the dense network of dark (gluconeogenic) cells form first, followed by a late appearance of light cells (Figures 6A and Video S1). This late appearance of light cells in the simulations comes from an inherent threshold effect included within the model, where the external build-up of the shared resource made by the dark cells is required. At a sufficient built-up concentration, this resource will trigger the switching of some cells to light cells. Light cells in turn will consume the resource, reducing the available amounts, thereby preventing other cells from switching to this new state. This threshold-effect therefore predicts a delayed, rapid emergence of light cells. If this threshold requirement is removed in the simulation (replaced by a rate of switching from dark to light that depends linearly on the amount of resource), the resultant colony remains small, and the organized pattern of cells in two states does not occur. This small colony size is largely due to low resource amounts to support the proliferation of the light cells, since there are insufficient dark cells remaining to produce the resource (Figures 6A, S5B, S5C, S5D and Video S5). In the model, the externally available amount of the resource builds-up, reaches the threshold (where cells switch to the light state), and then rapidly decreases, if the light cells also consume the resource (Figure 6B, upper panel). We therefore experimentally tested these model predictions, and first estimated the amounts of extracellular, free trehalose in the colony over time. Notably, trehalose amounts steadily increased over 4 days, and subsequently rapidly decreased (Figure 6B, lower panel). We next monitored the development of colonies over time, to determine when light cells emerge. Using just the bright-field image reconstruction of the colonies, during this time course, the intensity of dark cells steadily increased, and organized into the mesh-like network over 4 days (Figure 6C). However, the light cells appeared only after ∼4 days, and rapidly increased in number (Figure 6C). We more quantitatively estimated this, using strains expressing the gluconeogenic- or the PPP-reporter (Figure 6D). Notably, the increase in total fluorescence intensity due to the gluconeogenic-reporter in the colony (over time) was very linear over the first four days (r^2^=0.99). Contrastingly, the increase in the PPP reporter activity over the first five days was clearly non-linear, with very low signal intensity for the first three days, and then a rapid emergence of signal over days four and five (Figure 6E). This indicated a cooperative, switch like emergence of, and increase in these light cells. A useful biophysical measure of cooperativity (more commonly used for protein-ligand binding characteristics) is the Hill coefficient. We adopted the Hill equation, using the amount of PPP reporter fluorescence (instead of ligand-receptor binding), to estimate cooperativity in the system. Over the first five days the increase in PPP-reporter activity showed a Hill coefficient greater than 1, indicating a positively cooperative switch of cells to the light state (Figure 6E). This correlates perfectly with the build-up and rapid decrease in external trehalose (Figure 6B). In summary, data from model simulations and experiments show that initially the gluconeogenic cells increase in number, leading to release and build up the resource (trehalose) in the local environment. At a threshold concentration of trehalose, some cells switch to light state with high PPP activity. The further expansion of these light cells correlates with rapid consumption of the extracellular trehalose that sustains this state.

**Figure 6:**
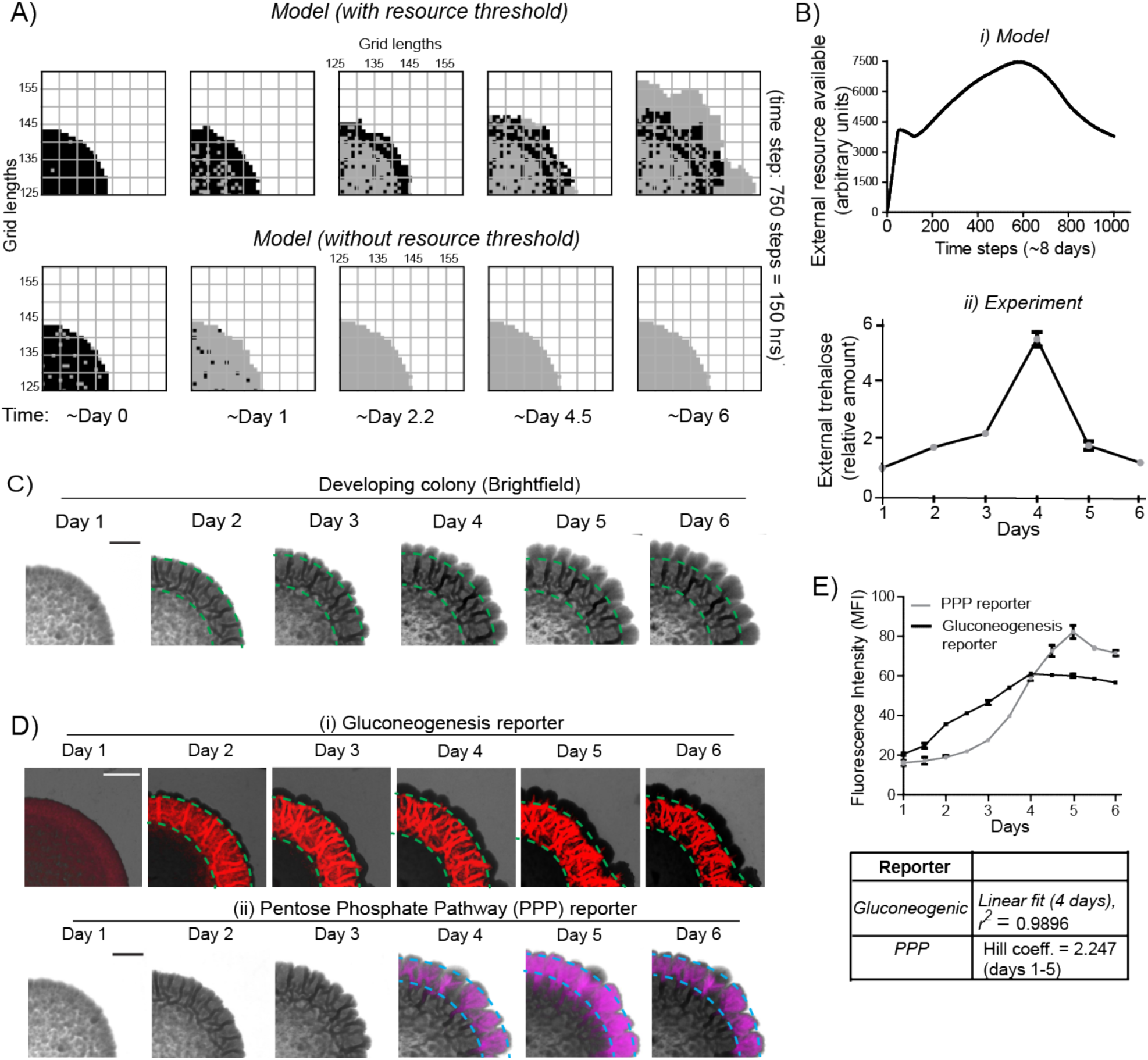
A resource threshold effect controls cooperative switching of cells to the light state. A) Simulation of colony development, based on the default model (which incorporates a resource threshold buildup, followed by consumption, switching to a light state, and expansion), compared to a model where the threshold amounts of the resource is removed. Note the final expansion size of the colony. Also see Figure S5 and Movie S5. B) i) Changes in the availability of the resource as the colony develops, based on the model. Ii) Extracellular amounts of trehalose measured from developing wild-type colonies. Data from three independent colonies. Note: in the model, in ∼3-4 days the resource is highest, and reduces sharply after that. In the experimentally obtained data, extracellular trehalose amounts are highest at ∼day 4, and then rapidly decreases over day 5. This correlates to when the light cells emerge and expand. C) A time-course of bright-field images of the developing wild-type colony, illustrating the distribution of dark cells, and the emergence and distribution of light cells. D) A time-course revealing fluorescence based estimation of the (i) reporter for gluconeogenic activity (dark cells), or (ii) the PPP activity reporter (light cells). Note the delayed, rapid appearance and increase in the PPP activity reporter. E) Quantification of the increase in the gluconeogenic reporter activity in the colony, and the PPP reporter activity (based on fluorescence intensity) within the colony. The increase in gluconeogenic reporter activity, when plotted, is linear, and saturates. The increase in PPP activity over the first 5 days is highly cooperative (as estimated using a Hill coefficient as a proxy for cooperativity), before saturating.

## Discussion

Collectively, we illustrate a simple schematic proposing how cells in phenotypically and metabolically distinct states spontaneously emerge and self-organize within a yeast colony. In low glucose conditions, cells begin in a uniform gluconeogenic state, which is the expected metabolic state in this nutrient condition. The gluconeogenic cells produce a resource (trehalose) that is externally available, and builds up to a threshold amount. At this threshold, if a cell takes up and consumes trehalose, this cell spontaneously switch to the complimentary metabolic state with high glycolytic and PPP activity (the light state) (Figure 7). Light cells remain in this metabolic state only as long as the resource (trehalose) is externally available. However, as trehalose is consumed by these cells, the available amount of external trehalose drops below the threshold, and the surrounding dark cells remain trapped in a gluconeogenic state, continuing to produce trehalose. A predictable fraction of cells, constrained spatially, will therefore remain in each metabolic state, resulting in specialized cell groups and division of metabolic labor. Thus, biochemically heterogeneous cell states can spontaneously emerge and spatially self-organize.

**Figure 7:**
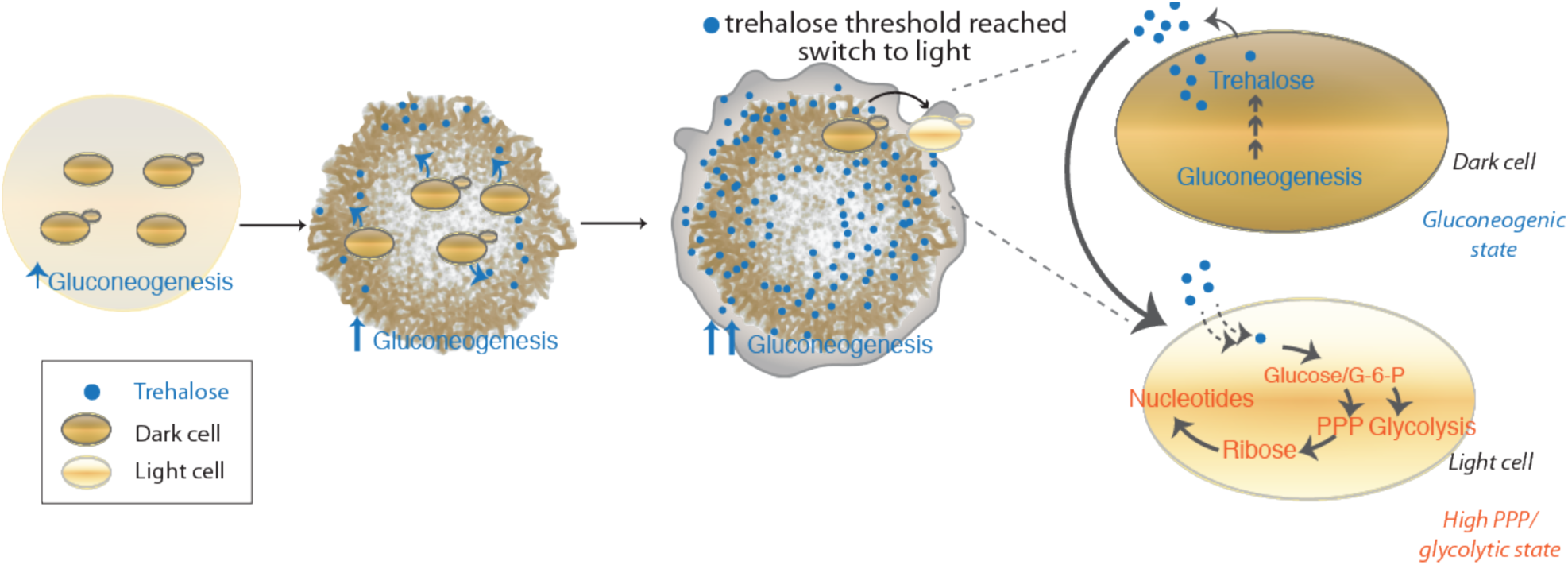
A simple model to explain emergent phenotypic heterogeneity and organization. Cells in low glucose perform gluconeogenesis, as required in low glucose medium. As gluconeogenic reserves build up, trehalose builds up in the extracellular environment. At a threshold concentration of trehalose, some cells switch to a high glycolytic, PPP state. This state depends upon the utilization of trehalose to fuel it. This utilization of trehalose by the light cells results in decreased external trehalose to below a threshold. This in turn restrains the other, remaining cells in a gluconeogenic state, where they continue to produce trehalose. This gives rise to the final, self-organized community, with specialization of function and division of labor.

Our study substantially advances descriptions of yeast ‘multicellularity’ from simple dimorphism, aggregated cells, or three-dimensional colony forms (Cáp et al., 2012; Koschwanez et al., 2011; Palková and Váchová, 2016; Ratcliff et al., 2012; Váchová and Palková, 2018; Veelders et al., 2010; Wloch-Salamon et al., 2017), to self-organized, phenotypically heterogeneous cell states exhibiting division of labor and metabolic interdependence. Strikingly, the nature of spatial patterning allied with division of labor that we observe in yeast is reminiscent of true multicellular systems (Newman, 2016; Niklas, 2014). Also, the cell states in these yeast colonies can be considered commensal, since trehalose is a necessary output of gluconeogenesis, and therefore a default, biochemically non-limiting output in dark cells. Since trehalose controls the emergence and maintenance of light cells in the complimentary metabolic state, it thus can be considered a resource benefiting the light state. Thus simple, metabolism-derived constraints are sufficient to determine how contrary biochemical states can spontaneously emerge and be supported, in conjunction with spatial structure. Such organization of cells into specialized, labor-divided communities expands on the role of reaction-diffusion systems (particularly activator-depleted substrate schemes) in controlling cellular patterning (Gierer and Meinhardt, 1972; Kondo and Miura, 2010; Newman, 2016), with a metabolic resource threshold being central to the emergence and stabilization of a new phenotype (Cai and Tu, 2011; Krishna and Laxman, 2018). A deeper dissection of what such constraints can permit will therefore advance our general understanding of how specialized cell states can emerge and be stabilized.

Metabolic cross-feeding is better understood in multi-species microbial communities, where this has been inferred largely using inter-species genomic comparisons (Ackermann, 2015; D’Souza et al., 2018; Goldford et al., 2018; Tyson et al., 2004). Further, metabolic sharing has typically been demonstrated using synthetically engineered systems where dependencies are created (Campbell et al., 2016; D’Souza et al., 2018; Mee et al., 2014; Pande et al., 2015; Wintermute and Silver, 2010). The spatial organizations of any such populations remain challenging to model. Biochemically identifying metabolites that are conclusively exchanged between cooperating cells remains difficult, and the significance of such putative metabolite exchange challenging to interpret (Ackermann, 2015; D’Souza et al., 2018). Finally, such studies have emphasized non-isogenic systems, where genetic changes stabilize different phenotypes, and auxotrophies define the nutrient sharing or cooperation (Ackermann, 2015). Contrastingly, here we directly identify a produced metabolic resource, and demonstrate how its availability and differential utilization can control the emergence of cells in opposing metabolic states, in a clonal population. This also explains the spontaneous spatial organization into phenotypically distinct cell groups. Thus, our study also goes beyond stochastic gene expression (Ackermann, 2015; Balázsi et al., 2011; Blake et al., 2003) to explain how phenotypic heterogeneity and specialization can emerge in clonal populations. By considering these metabolism-derived rules, and thereby manipulating available metabolic resources, it may be viable to program the formation, structure or phenotypic composition of isogenic cell populations. Collectively, such simple physico-chemical constraints can advance our understanding of how isogenic cells can self-organize into specialized, labor-divided groups, as a first step towards multicellularity.

## Experimental Procedures

### Yeast strains and growth media

The prototrophic sigma 1278b strain (referred to as wild-type or WT) was used in all experiments. Strains with gene deletions or chromosomally tagged proteins (at the C-terminus) were generated as described (Longtine et al., 1998). Strains used in this study are listed in Table 1. The growth medium used in this study is rich medium (1% yeast extract, 2% peptone and 2% glucose or 0.1% glucose).

### Colony spotting assay

All strains were grown overnight at 30°C in either rich medium or minimal medium. 5 microliters of the overnight cultures were spotted on rich medium (low glucose) (1% yeast extract, 2% peptone, 0.1% glucose and 2% agar). Plates were incubated at 30°C for 7 days unless mentioned otherwise.

### Colony imaging

For observing colony morphology, colonies were imaged using SZX-16 stereo microscope (Olympus) wherein the light source was above the colony. Bright-field imaging of 7-day old colonies were done using SZX-16 stereo microscope (Olympus) wherein the light source was below the colony. Epifluorescence microscopy imaging of 7-day old gluconeogenesis reporter colonies (pPCK1-mCherry), pentose phosphate pathway (PPP) reporter colonies (pTKL1-mCherry) and *HXK1* reporter colonies (pHXK1-mCherry) were imaged using the red filter (excitation of 587 nm, emission of 610 nm) of SZX-16 stereo microscope (Olympus). Similar protocol was followed for imaging 1-day to 6-day old colonies.

### Analysis of fluorescent cell populations in reporter strain colonies

Light cells and dark cells isolated from 7-day old wild-type colonies harboring either the gluconeogenesis reporter, PPP reporter or the *HXK1* reporter were re-suspended in 1 ml of water. The percentage of fluorescent cells were determined by running the samples through a flow cytometer, and counting the total number of mCherry positive cells in a total of 1 million cells. Light cells and dark cells isolated from wild-type colonies without the fluorescent reporter were used as control.

### Biochemical estimation of trehalose/glycogen levels

Trehalose and glycogen from yeast samples were quantified as described previously, with minor modifications (Shi et al., 2010b). 10 OD_600_ of light cells and dark cells from 7-day old wild-type colonies (rich medium, 0.1% glucose) were collected. After re-suspension in water, 0.5 ml of cell suspension was transferred to 4 tubes (2 tubes for glycogen assay and the other 2 tubes for trehalose assay). When sample collections were complete, cell samples (in 0.25 M sodium carbonate) were boiled at 95–98°C for 4 hr, and then 0.15 ml of 1 M acetic acid and 0.6 ml of 0.2 M sodium acetate were added into each sample. Each sample was incubated overnight with 1 U/ml amyloglucosidase (Sigma-Aldrich) rotating at 57°C for the glycogen assay, or 0.025 U/ml trehalase (Sigma-Aldrich) at 37°C for the trehalose assay. Samples were then assayed for glucose using a glucose assay kit (Sigma-Aldrich). Glucose assays were done using a 96-well plate format. Samples were added into each well with appropriate dilution within the dynamic range of the assay (20–80 µg/ml glucose). The total volume of sample (with or without dilution) in each well was 40 microliters. The plate was pre-incubated at 37°C for 5 min, and then 80 µl of the assay reagent from the kit was added into each well to start the colorimetric reaction. After 30 min of incubation at 37°C, 80 microliters of 12 N sulfuric acid was added to stop the reaction. Absorbance at 540 nm was determined to assess the quantity of glucose liberated from either glycogen or trehalose. For measurement of extracellular trehalose measurement, single wild-type colony (1-day to 7-day old colony) was re-suspended in 100 microliters of water and centrifuged at 20000g for 5 min. Supernatant was collected and buffered to a pH of 5.4 (optimal for trehalase activity) using sodium acetate buffer (pH 5.0). 0.025 U/ml trehalase (Sigma-Aldrich) was added and samples were incubated at 37°C overnight. Glucose concentration was estimated as described earlier.

### Neutral trehalase activity assay

Neutral trehalase activity assay was performed as described earlier with the following modifications (De Virgilio et al., 1991). Briefly, 1 OD_600_ of light cells and dark cells isolated from 7-day old wild-type colonies (rich medium, 0.1% glucose) were washed twice with ice-cold water. For permeabilization, cells were re-suspended in tubes containing equal volume of 1% Triton-X in assay buffer (200mM tricine buffer (Na^+^) (pH 7.0)) and immediately frozen in liquid nitrogen. After thawing (1-4 min at 30°C), the cells were centrifuged (2 min at 12000g), washed twice with 1ml of ice-cold assay buffer and immediately used for the trehalase assay. Trehalase assay was performed in 50mM tricine buffer (Na^+^) (pH 7.0), 0.1 M trehalose, 2mM manganese chloride (MnCl_2_) and the Triton X-100 permeabilized cells in a total volume of 400 microliters. After incubation for 30 min at 30°C, the reaction was stopped in a boiling water bath for 3 min. Glucose concentration in the supernatant was determined using the glucose assay kit (Sigma-Aldrich).

### Western blot analysis

Approximately ten OD_600_ cells were collected from respective cultures, pelleted and flash frozen in liquid nitrogen until further use. The cells were re-suspended in 400 microliters of 10% trichloroacetic acid (TCA) and lysed by bead-beating three times: 30 sec of beating and then 1 min of cooling on ice. The precipitates were collected by centrifugation, re-suspended in 400 microliters of SDS-glycerol buffer (7.3% SDS, 29.1% glycerol and 83.3 mM tris base) and heated at 100°C for 10 min. The supernatant after centrifugation was treated as the crude extract. Protein concentrations from extracts were estimated using bicinchoninic acid assay (Thermo Scientific). Equal amounts of samples were resolved on 4 to 12% bis-tris gels (Invitrogen). Western blots were developed using the antibodies against the respective tags. We used the following primary antibody: 538 monoclonal FLAG M2 (Sigma-Aldrich). Horseradish peroxidase–conjugated secondary antibody (anti-mouse) was obtained from Sigma-Aldrich. For Western blotting, standard enhanced chemiluminescence reagents (GE Healthcare) were used.

### Metabolite extractions and measurements by LC-MS/MS

Light cells and dark cells isolated from wild-type colonies grown in different media were rapidly harvested and metabolites were extracted as described earlier (Walvekar et al., 2018). Metabolites were measured using LC-MS/MS method as described earlier (Walvekar et al., 2018). Standards were used for developing multiple reaction monitoring (MRM) methods on Sciex QTRAP 6500. Metabolites were separated using a Synergi 4µ Fusion-RP 80A column (100 × 4.6 mm, Phenomenex) on Agilent’s 1290 infinity series UHPLC system coupled to the mass spectrometer. For positive polarity mode, buffers used for separation were-buffer A: 99.9% H_2_O/0.1% formic acid and buffer B: 99.9% methanol/0.1% formic acid (Column temperature, 40°C; Flow rate, 0.4 ml/min; T = 0 min, 0% B; T = 3 min, 5% B; T = 10 min, 60% B; T = 11 min, 95% B; T = 14 min, 95% B; T = 15 min, 5% B; T = 16 min, 0% B; T = 21 min, stop). For negative polarity mode, buffers used for separation were-buffer A: 5 mM ammonium acetate in H_2_O and buffer B: 100% acetonitrile (Column temperature, 25°C; Flow rate: 0.4 ml/min; T = 0 min, 0% B; T = 3 min, 5% B; T = 10 min, 60% B; T = 11 min, 95% B; T = 14 min, 95% B; T = 15 min, 5% B; T = 16 min, 0% B; T = 21 min, stop). The area under each peak was calculated using AB SCIEX MultiQuant software 3.0.1.

### ^15^N- and ^13^C-based metabolite labelling experiments

For detecting ^15^N label incorporation in nucleotides, ^15^N Ammonium sulfate (Sigma-Aldrich) and ^15^N Aspartate (Cambridge Isotope Laboratories) with all nitrogens labeled were used. For ^13^C-labeling experiment, ^13^C Trehalose with all carbons labeled (Cambridge Isotope Laboratories) was used. All the parent/product masses measured are enlisted in Table 2. For all the nucleotide measurements, release of the nitrogen base was monitored in positive polarity mode. For all sugar phosphates, the phosphate release was monitored in negative polarity mode. The HPLC and MS/MS protocol was similar to those explained above.

## Building and implementing the model

### Components

The model consists of (i) a population of light and dark cells, and (ii) a shared metabolic resource that is produced by, and is accessible to the cells. Therefore, the dynamic processes involved can be broadly divided into those pertaining to the cells of the colony and those pertaining to the shared resource. The cells and resource occupy a 2-D square grid, which represents the surface of an agar plate. If one takes each grid length to correspond to 50µm in real space, then, given the average size of a yeast cell at 5µm, a single grid location can be imagined to contain upto 100 cells, which we term “cell blocks”. We coarse-grain the model such that each location is either empty, occupied by light cell block, or a dark cell block. That is, we ignore the possibility that cell blocks might be mixed. This is simply for computational ease. A more detailed model consisting of smaller grid lengths such that each location could hold at most a single cell would exhibit the same behavior as the coarse-grained one, but would require much larger grid sizes and longer computational times in order to simulate realistic sized colonies. With the coarse-graining, our simulations use a 250 × 250 grid. Each grid location also contains saturating amounts of amino acids, as well as a certain level of the shared metabolic resource. If a location has a cell block, that block also has internal levels of the amino acids and the resource, which may be different from the external level in that location.

### Initial state of the grid

We start with an approximately circular colony 20 grid lengths in radius (covering 1257 grid locations) in the center of the 250 × 250 grid. 95-99% of these 1257 cell blocks are in the dark state, while 1-5% are in the light state, distributed randomly in the colony. The concentration or level of the shared resource is set to zero at every location. However, at all times, we assume the presence, throughout the grid, of saturating amounts of amino acids that are required for the (slow) growth of the dark cells.

### Dynamics of the model

The grid is updated at discrete time steps. Each time step corresponds to 12 min in real time, and all simulations are run for 750 time steps, i.e., 150 hours of real time (∼ 6 days). In each time step, we first go over every cell block to implement the following processes:

If a block at location (x,y) is dark, then:

1. If the resource level at (x,y) is above a certain threshold **S** = 3.0 units of resource, then the cells in the block switch to being light cells with a probability **p** = 0.5
2. If the block is still dark, then add **R** = 0.07 units to the resource level at (x,y).
3. Consume (internalize) **C** = 0.05 units of amino acids (present in saturating amounts at all locations)
4. If the internal amino acid level is greater than or equal to 1.0, the dark block can divide with a probability **g**_**d**_ = 0.01.
5. If the block can divide, then check if there’s an empty location in the immediate neighborhood. The immediate neighborhood is the set of locations {(x-1,y), (x+1,y), (x,y-1), (x,y+1)}.
6. If there’s at least one empty space, preferably divide into an empty location which has more occupied neighbors. After division, the two daughter blocks are each assigned half the internal amino acid reserves of the original mother block.

If a block at location (x,y) is light, then :

1. If the resource level at (x,y) is greater than or equal to **C** = 0.05 units, consume (internalize) all of it.
2. If the internal resource level is greater than or equal to 1.0, the dark block can divide with a probability **g**_**l**_ = 0.04.
3. If the block can divide, then check if there’s an empty location in the immediate neighborhood. The immediate neighborhood is the set of locations {(x-1,y),(x+1,y),(x,y-1),x,y+1)}.
4. If there’s at least one empty space, preferably divide into an empty location that has more occupied neighbors. After division, the two daughter blocks are each assigned half the internal resource reserves of the original mother block.

(The above set of rules and parameters is for simulating the wild-type colony. For the variations highlighted in the main text (Figure 3E and 6A, bottom row), see the ‘Variants of the wild-type model’ section below.)

These processes implement growth of cells, as well as production and consumption of amino acids and the shared metabolic resource. Subsequent to this, in each time step, we allow diffusion of the resource levels across the grid (the “external” level at the location, not the internal levels in cell blocks), using a numerical scheme called Forward Time Central Space (FTCS). Say that the value of the resource at time t and location (x,y) is given by 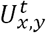. The FTCS scheme updates the value simultaneously at all locations using the following formula:

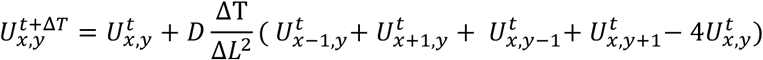

where *ΔT* is the time step and *ΔL* is the space step, or grid length, and *D* is the diffusion constant for the resource.

### Model parameters

1) The parameters of the model are shown in Table 3. Time and length units are chosen such that each time step is one unit of time, and each grid length is one unit length. With these choice of units, the growth parameters for light and dark cells, respectively, are g_l_ = 0.04, g_d_ = 0.01. These were chosen so as to reflect the relative rates of diffusion and division. Light cells were observed to grow faster than the dark cells, so their respective growth parameters are set accordingly.

**Table 3.** Parameters of the model for the wild-type case.

| Parameter | Notation | Value |
| --- | --- | --- |
| Growth rate (dark cell block) | $g_d$ | 0.01 /T |
| Growth rate (light cell block) | $g_l$ | 0.04 /T |
| Switching threshold | $S$ | 3.0 units |
| Resource produced by each dark cell block | $R$ | 0.07 units/T |
| Resource or amino acids consumed per cell block (light or dark) | $C$ | 0.05 units/T |
| Minimum resource or amino acid reserve needed for division (light or dark) | -- | 1.0 unit |
| Chance to switch to light cell if threshold reached | $P$ | 0.5 /T |
| Diffusion constant of the resource | $D$ | 0.24 L<sup>2</sup>/T |

2) The switching threshold parameter (S = 3.0) was chosen to account for a delay in the switching of dark cells to another metabolic pathway via nutrient sensing as well as to give a reproducible facsimile of the experimental colonies.

3) The shared resource production value was chosen to be 7% (R = 0.07) of the minimum required to divide. In each time step, every block of dark cells adds this amount to the resource grid. This was chosen as a default value, which gave a reproducible facsimile of the experimental colonies. Other values were tried and their effect is seen in Figure S4.

4) All cells consumed a small level of metabolites (the shared resource or amino acids) in each time step. This value was chosen to be 5% of the minimum required for division (C = 0.05). This gave division times that approximately matched the division times observed experimentally.

5) The switching probability (p = 0.5) was chosen to add an element of stochasticity. So even if the threshold resource conditions are met, dark cells have a 50% chance to switch to light cells in that time step.

6) The choice of the diffusion constant (D = 0.24) is limited by the numerical stability of the FTCS scheme, which allows only a maximum value of D = 0.25. In real time and length units, this corresponds to a diffusion constant D_eff_ of 8.7 × 10^−13^ m^2^/s. D_eff_ is an order of magnitude smaller than the diffusion constant for sugars like glucose and sucrose in water *(57)*. Since the agar used for the experiments is mostly water, the diffusion constants in water can be considered as a good reference point.

### Variants of the wild-type model in different figures

Figures 3E, Figure 6 (bottom row) and Figure S5 showcase some of the final colonies generated by the simulations when the rules described above are varied. The following changes were made to the rules/parameters to generate these.

3E(i): No sharing: Set R = 0.

3E(ii): No switching from dark to light state: Set p = 0.

3E(iii): ‘Reverse’ sharing: Set R = 0. When a cell block is light it adds R’ = 0.07 to the amino acid grid at the same location.

6(bottom row): No resource thresholding: Set S = 0.

S5: Linear switching: Set S = 0. The probability of switching from dark to light state, *p*, is now a linear function of the locally available resource with a maximum value of 1.0. That is, 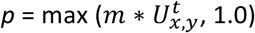, where *m* is a parameter that sets the slope of this linear relationship.

## Author contributions

SV designed and performed experiments, analyzed and interpreted data, and contributed to writing the paper. AW designed and performed experiments, analyzed and interpreted data. VS designed and implemented the mathematical model, carried out simulations, and contributed to writing the paper. SK designed the mathematical model, helped validate it, and contributed to writing the paper. SL conceived the project, designed experiments, analyzed and interpreted data, supervised the project and wrote the paper.

## Data and materials availability

all strains used in this study are available on request. The model simulation code is available via GitHub ref: https://github.com/vaibhhav/yeastmetabcolony.

## Supplementary Figures

**Figure S1:**
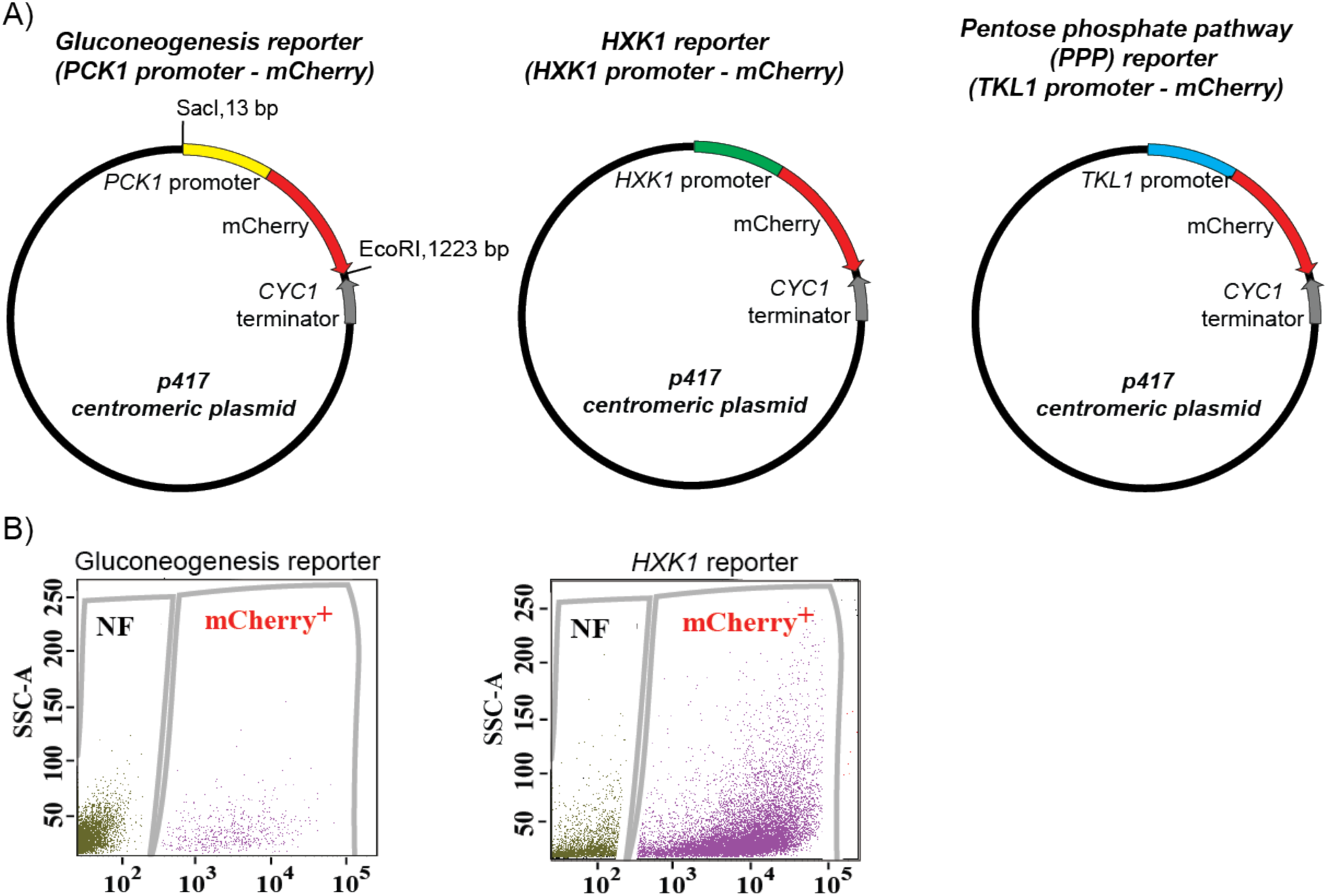
Gluconeogenesis activity is spatially restricted. (A) Plasmid maps of the different reporters used in this study. Gene cassette containing the specific promoter and mCherry was cloned into the p417 centromeric plasmid using the Sacl and EcoRl restriction sites. (B) Scatter plot depicting distribution of mCherry positive light cells isolated from wild-type colonies with the gluconeogenesis or HXK1 reporter. SSC-A represents side scatter measurement and NF represents non-fluorescent population.

**Figure S2:**
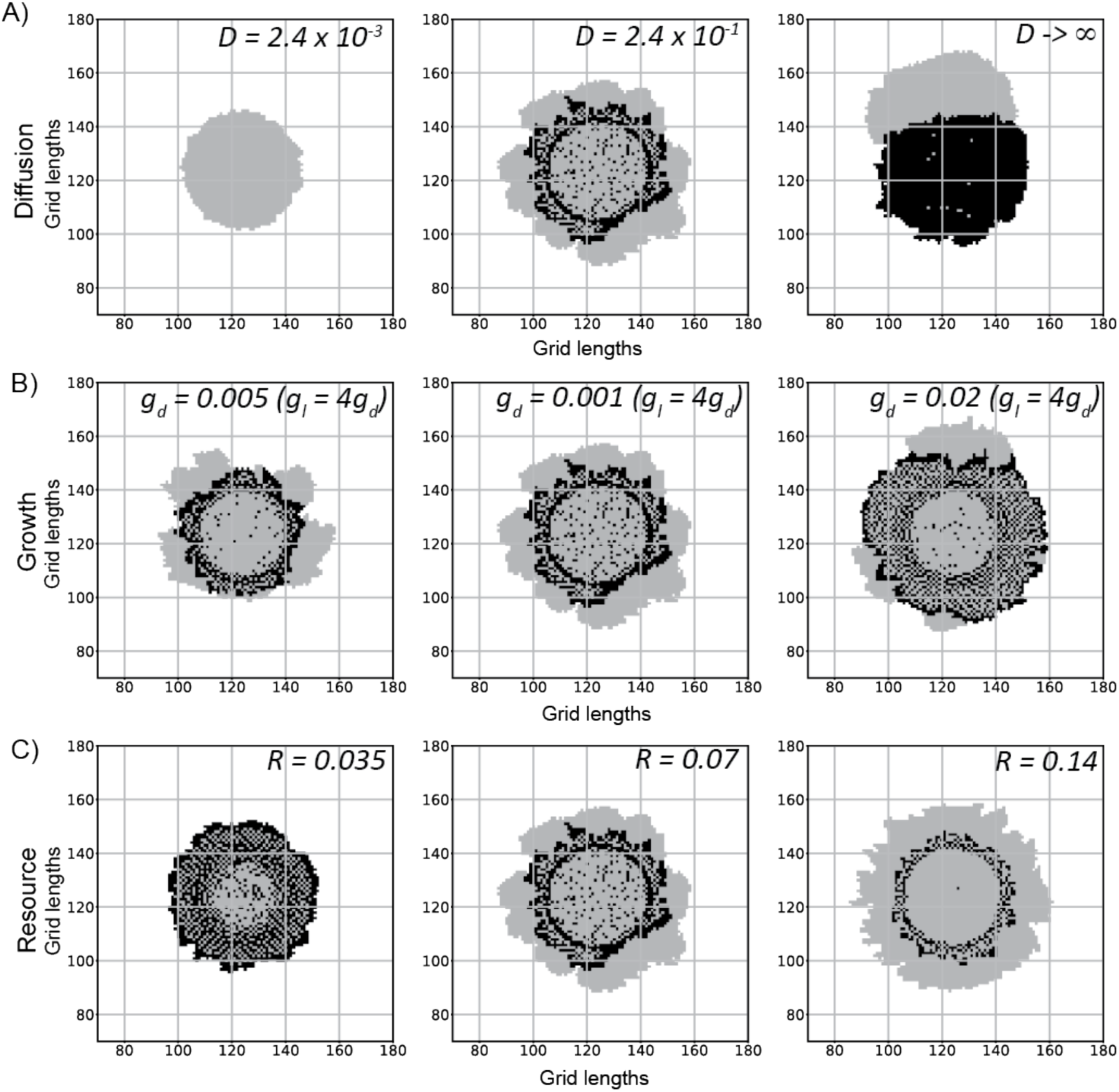
Effects on the colony as we change single parameters used in the model. (A) Changing the diffusion constant (D [L^2^T^−1^])of the resource in the medium. The diffusion constant used to simulate the left panel colony is 100 times smaller than the default colony (middle panel). The right panel colony was simulated with a very large effective diffusion constant. To get this fast diffusion, the total available resource was uniformly redistributed over the grid between every two time steps. Thus, every grid location had the same level of resource before the cells consume or produce it, as if the resource diffused very quickly over the grid. (B) Changing the growth rate of the cells. g_d_ and g_i_ are growth rate parameters of the dark and light cells respectively. The light cells grow 4 times faster than the dark cells in all cases. The middle panel colony is the default colony. g_d_ = 0.01. The one on the left has half the growth rate and the one on the right has twice the growth rate in comparison. (C) Changing the resource (R) produced by the dark cells. The light cells consume resource at a fixed rate if available. The middle panel has R = 0.07. The left colony simulation has half the produced value and the right colony simulation has double the produced value. All colony simulations were started with 1257 grids (each containing ∼100 cells), of which 99% were dark cells, and were run for 750 time steps.

**Figure S3:**
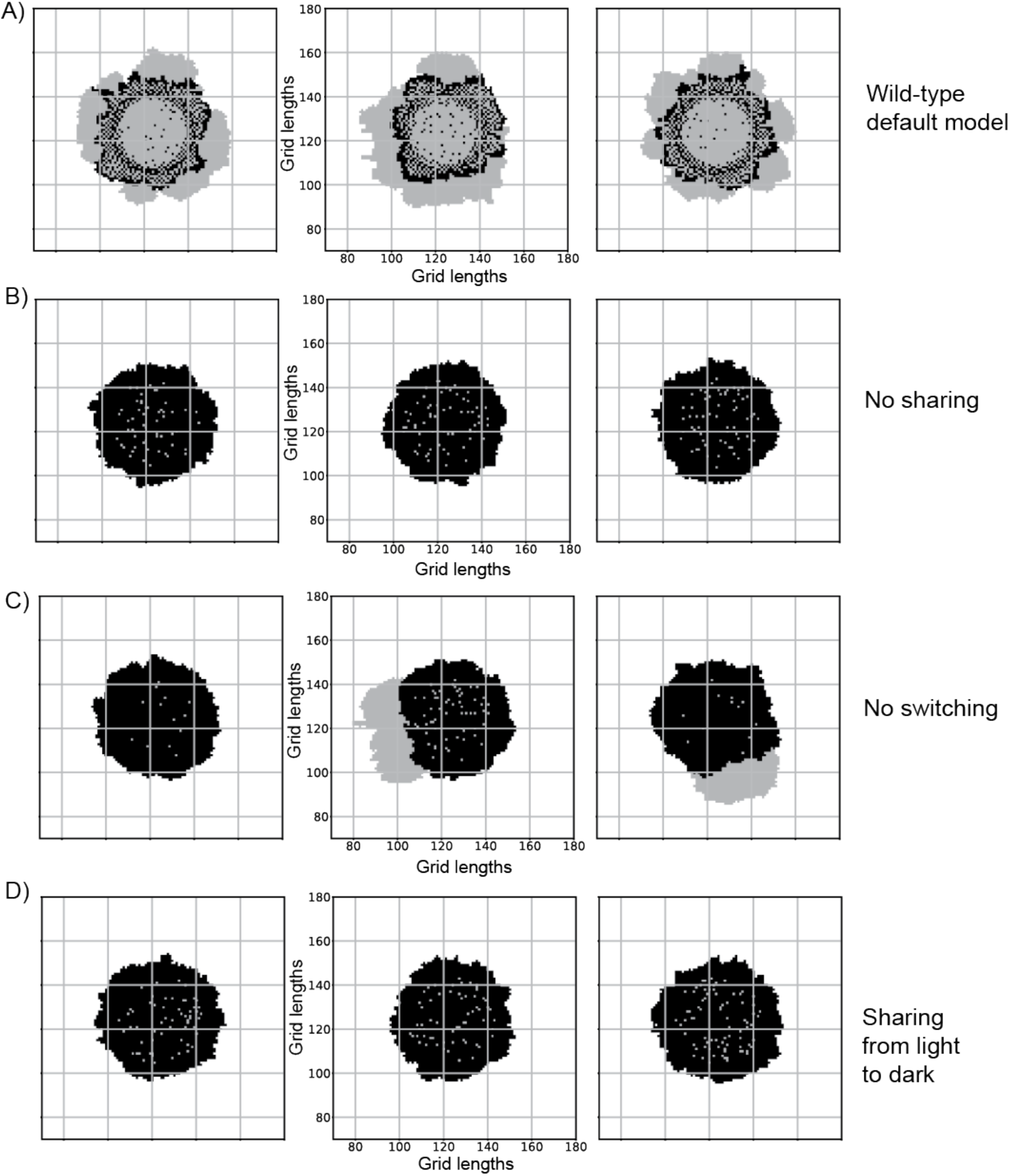
Reproducibility of the model, under different scenarios. Since the colony simulations have elements of stochasticity present in the model, presented are a few replicate colonies from independent simulation. These showcase crucial and reproducible aspects of the different conditions explored in figure 4D and figure 4E. (A) 3 wild-type colonies generated with the default parameter set. The common features are a circular centre with mostly light cells, an annular region with dark cells and a few light cells, and the colony periphery with mostly light cells. (B) 3 replicate colonies with the “no sharing” condition. The dark cells do not produce any shared resource. The final colonies have mostly dark cells as the number of light cells doesn’t change. (C) 3 replicate colonies with the “no switching” condition, if there are a few light cells at the periphery, they can consume the shared resource produced by the dark cells and grow into the empty space. However, the centre of the colonies has mostly dark cells. (D) 3 replicate colonies with the “wrong sharing” condition (sharing of a resource made by light cells, taken up by dark cells), which are similar to the “no sharing” condition.

**Figure S4:**
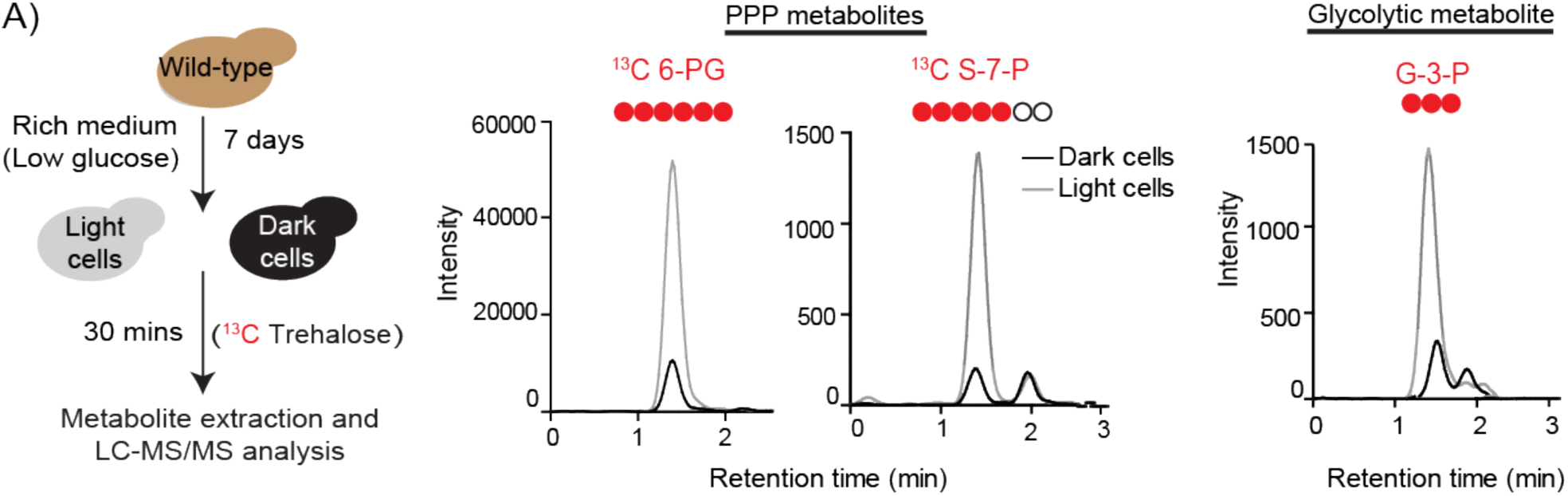
Comparative breakdown of labeled trehalose by distinct cells in a colony. (A) Trehalose breakdown by light cells and dark cells were monitored by isolating light and dark cells from a ∼5 to 6-day old wild-type colony and incubating them briefly (30 min) with labeled ^13^C Trehalose. Post incubation, metabolites from light and dark cells were extracted and analyzed for the pentose phosphate pathway (PPP) intermediates ^13^C 6-phosphogluconate (^13^C 6-PG),^13^C Sedoheptulose-7-phosphate (^13^C S-7-P) and glycolytic intermediate ^13^C Glyceraldehyde-3-phosphate (^13^C G-3-P) using LC-MS/MS.

**Figure S5:**
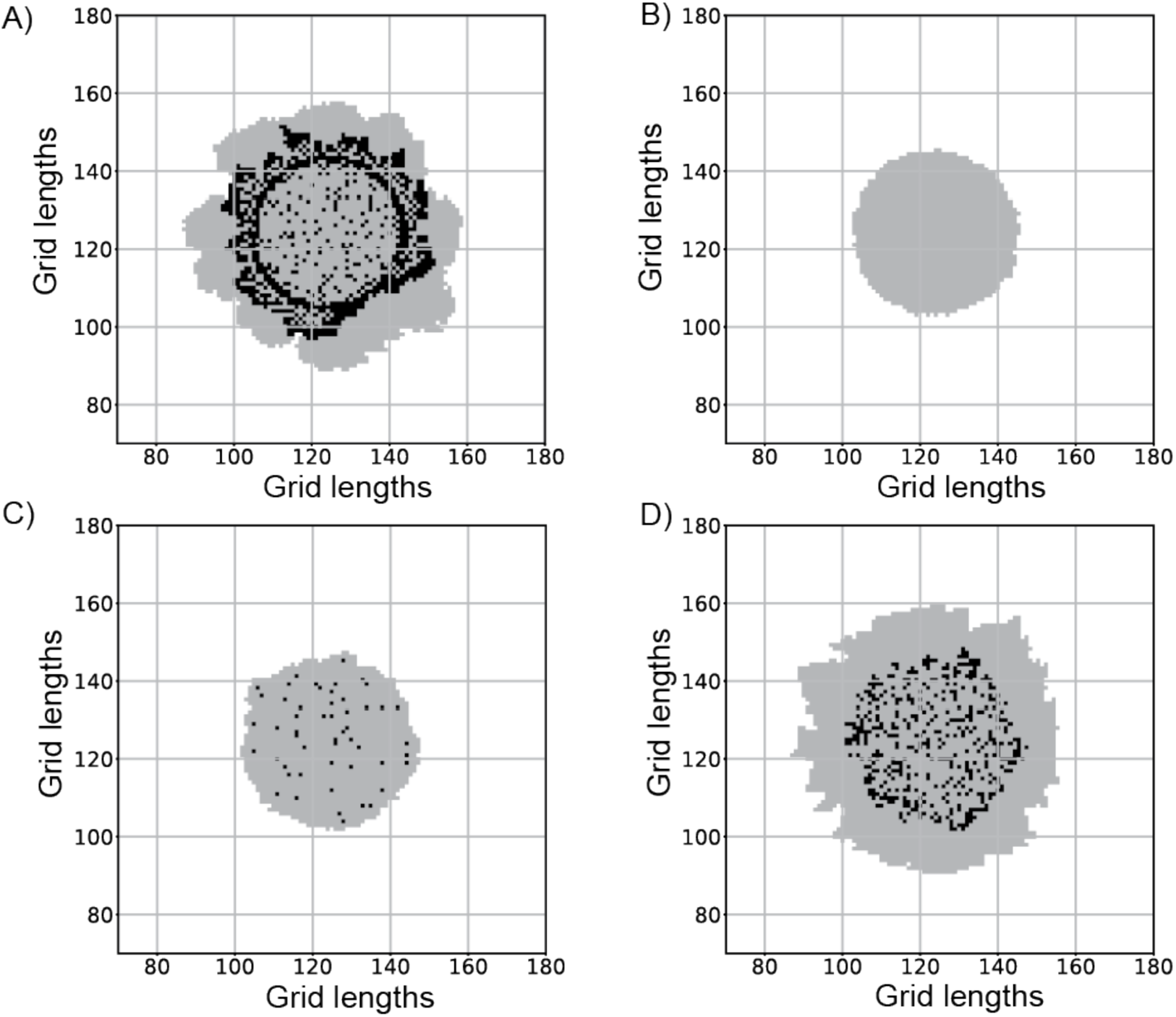
Colonies generated using different switching rules. (A) Wild-type colony with a resource threshold switching rule. In every time step, the algorithm checks if the shared resource levels at the locations of dark cells are more than a threshold value of 3 units. If so, there is a 50% chance of those cells switching to light cells. (B) A colony generated without the need for a resource threshold level. Dark cells switch by random chance. Light cells cannot switch to dark. (C) & (D) Colonies generated using a linear switching rule. The probability of switching increases linearly with the locally available resource levels till a maximum value of 1.0. The probability of switching is of the form p = mU) (if mU less than 1.0) and p = 1.0 otherwise. Here U is the resource level at the location of the dark cells and m is the slope of the linear relationship. In panel (C) m = 1. In panel (D) m = 0.01.

